# Cuproptosis is correlated with clinical status, tumor immune microenvironment and immunotherapy in colorectal cancer: a multi-omic analysis

**DOI:** 10.1101/2022.09.12.507555

**Authors:** Yanfei Shao, Xiaodong Fan, Xiao Yang, Shuchun Li, Ling Huang, Xueliang Zhou, Sen Zhang, Minhua Zheng, Jing Sun

**Affiliations:** Department of General Surgery, Ruijin Hospital, Shanghai Jiao Tong University School of Medicine, Shanghai, China; Shanghai Minimally Invasive Surgery Center, Ruijin Hospital, Shanghai Jiao Tong University School of Medicine, Shanghai, China; Shanghai Institute of Digestive Surgery, Ruijin Hospital, Shanghai Jiao Tong University School of Medicine, Shanghai, China

**Keywords:** Colorectal Cancer, Cuproptosis, Molecular Phenotypes, CuproScore, Immunotherapy

## Abstract

Copper, like double-edged sword, either too little or too much can lead to cell death. Cuproptosis, a novel identified cell death form induced by copper, is characterized by aggregation of lipoylated mitochondrial enzymes and the destabilization of Fe–S cluster proteins. However, the function and potential clinical value of cuproptosis in colorectal cancer remains largely unknown. In this study, 16 cuproptosis-related molecules (CPRMs) were identified and analyzed by transcriptomics, genomics, and single-cell transcriptome profiling from multiple databases. We established two cuproptosis-related molecular phenotypes (CMP1 and 2) to distinguish CRC individuals, in which there were significantly differences in prognosis, biological function, somatic mutation frequency, immune cell infiltration in CRC individuals. A novel cuproptosis-related scoring system (CuproScore) was also constructed to predict the prognosis of CRC individuals, TME and the response to immunotherapy. Of notion, the value of CuproScore was also confirmed in our transcriptome cohort, demonstrating that CRC individuals in the high CuproScore group tended to have higher immune cell infiltrations and higher immune checkpoint expression. We also checked and analyzed the expression and clinical significance of 16 CPRMs in CRC cell lines and CRC tissues. This study indicated that cuproptosis and CPRMs played significant role in CRC progression and in modeling the TME. Inducing cuproptosis may be a useful tool for tumor therapy in the future.

## INTRODUCTION

Colorectal cancer (CRC) is one of the most common malignancies in the world, ranking second for incidence (more than 1,900,000 cases) and third for mortality (more than 930,000 cases) worldwide in 2020^1^. Recently, with the development of comprehensive treatments in CRC, including curative surgery, neoadjuvant chemoradiotherapy, systemic chemotherapy, targeted therapy, the prognosis of CRC individuals has been significantly improved^2, 3^. Furthermore, tumor immunotherapy, especially immune checkpoint inhibitors (ICIs), has shown exciting clinical success in various solid tumor treatments^4^. Unfortunately, the prognosis for individuals diagnosed with CRC at an advanced stage remains poor, and only a few CRC individuals with microsatellite instability high, approximately 4-5% of mCRC population, could be benefiting from ICIs^5^. Thus, more reliable and accurate biomarkers and potential therapeutic targets should be explored to improve the diagnosis, prognosis, and therapeutic effect of CRC individuals.

Cell death is an inevitable and complicated process in both normal and tumor cells. Different from the other well-known types of regulated cell death (RCD), such as apoptosis^6^, autophagy^7^, pyroptosis^8^, and ferroptosis^9^, cuproptosis is a copper-induced cell death reported in a recent study by Tsvetkov et al^10^, characterized by excess intracellular copper, lipoylated mitochondrial enzymes accumulation and Fe-S cluster proteins loss. Besides, cuproptosis has also been considered to have a close association with the tumor metabolism, such as the tricarboxylic acid (TCA) cycle. The metabolic reprogramming occurs throughout the process of malignant transformation, to meet the demands of the tumor microenvironment (TME) and unique tumor cell growth^11, 12^. In addition, the process of metabolic reprogramming also plays central role in CRC to enable tumor proliferation in the hypoxia condition and the tumor immune escape^13–15^. Therefore, cuproptosis is likely to be involved in tumorigenesis and TME in CRC. However, there is few study deep exploring the role of cuproptosis in CRC, and whether cuproptosis-related molecules (CPRMs) could regulate the pathogenesis of CRC remains largely unknown.

Therefore, to get a better understanding of cuproptosis in the tumorigenesis and clinicopathological characteristics in CRC individuals, we presented a multi-omic analysis of the CPRMs in CRC, including the clinical status, molecular functions, and tumor immune microenvironment. Our results reveal that cuproptosis plays a significant role in the tumorigenesis of CRC with potential influence on the tumor microenvironment. CuproScore is a potentially powerful scoring system for predicting prognosis and the response to immunotherapies in CRC individuals. These findings will help understand cuproptosis in regulating CRC profiles and provide new directions for therapeutic intervention in the future.

## Materials and Methods

### Collection of Data

The pan-cancer RNA sequencing (RNA-seq), corresponding clinical characteristics, somatic mutation, and copy number variation data in the Cancer Genome Atlas (TCGA) datasets were downloaded from the Genomic Data Commons (GDC, https://portal.gdc.cancer.gov/). The RNA-seq data of the normal tissues in the Genotype-Tissue Expression (GTEx) were downloaded from the University of California Santa Cruz (UCSC, https://xenabrowser.net/datapages/). All the RNA-seq data were converted from count values to Transcripts Per Kilobase of exon model per Million mapped reads (TPM) values. Microsatellite Instability(MSI) data of TCGA colorectal cancer patients^16^ was obtained by R package ‘TCGAbiolinks’ ^17^. The Consensus Molecular Subtypes (CMS)^18^ of CRC cohort in TCGA COAD and READ datasets were calculated by R package ‘CMScaller’ (https://github.com/Lothelab/CMScaller) ^19^. Besides, information of metastatic or recurrent CRC patients who received 5-Fluorouracil (5-Fu), FOLFOX6, FOLFIRI, Cetuximab, PD-L1, or CTLA-4 treatments were respectively obtained in the Gene-Expression Omnibus (GEO) database (GSE62080, GSE19862, GSE19860, GSE108277, and GSE81005, https://www.ncbi.nlm.nih.gov/geo/) and all the RNA-seq and clinical characteristics were obtained by R package ‘GEOquery’ ^20^. In addition, the single-cell RNA-seq data of the GSE132465 cohort was also downloaded from the GEO database (https://www.ncbi.nlm.nih.gov/). The RNA-seq data of the CRC cell lines were downloaded from the Cancer Cell Line Encyclopedia (CCLE, https://sites.broadinstitute.org/ccle/)^21^. The accessible immunohistochemical images of CRC individuals were obtained from the Human Protein Atlas (HPA, https://www.proteinatlas.org/) database^22^. Furthermore, the complete RNA-seq data and corresponding clinical characteristics of the IMvigor210 cohort^23^ (the research cohort of anti-PD-L1 immunotherapy in bladder cancer) and Liu’s cohort^24^ (the research cohort of anti-PD-1 immunotherapy in melanoma) were obtained from the link (IMvigor210 cohort, http://research-pub.gene.com/IMvigor210CoreBiologies/; Liu’s cohort, https://github.com/vanallenlab/schadendorf-pd1).

### Generation of Cuproptosis-associated Molecules

16 cuproptosis-associated molecules (FDX1, LIPT1, LIAS, DLD, DBT, GCSH, DLST, DLAT, PDHA1, PDHB, SLC31A1, ATP7A, ATP7B CsDKN2A, GLS, and MTF1) were obtained from the previous study^10^, including components of the lipoic acid pathway, components of the pyruvate dehydrogenase (PDH) complex and copper transporters. The biological information of these genes was shown in Supplementary Table S1.

### Analysis of Cuproptosis-related Genes in Pan-cancer

For differential expression analysis, the R package ‘DESeq2’^25^ was used to compare the cuproptosis-related gene expression levels between normal and tumor tissue among pan-cancer. The prognostic values of cuproptosis-related genes were determined by univariable Cox regression among CRC and other types of cancer. The R package ‘maftools’ was used to draw a waterfall chart of cuproptosis-related gene somatic mutation and estimate the Tumor Mutation Burden (TMB) of CRC individuals in TCGA datasets^26^. The correlation between cuproptosis-related gene expression and copy number in pan-cancer individuals was calculated by Spearman rank correlation analysis. The gain or loss of copy numbers of cuproptosis-related genes were determined by the total number of genes with copy numbers changing at the focal and arm levels. The correlation of cuproptosis-related gene expression levels was calculated by the R packages “igraph” and “reshape2”, and the protein-protein interaction (PPI) network was conducted to reveal the interaction of proteins among the proteins coding between the 16 cuproptosis-related genes by the STRING database (http://www.string-db.org/). Molecular pathways involved in these cuproptosis-related genes were estimated by the R package ‘ClusterProfiler’^27^ for the gene ontology (GO) and Kyoto Encyclopedia of Genes and Genomes (KEGG) analysis.

### Identification of the enrichment scores of gene features

The cuproptosis-related pathway was identified based on the 16 cuproptosis-related genes. The cuproptosis-related pathway enrichment scores of each sample were evaluated by R package ‘GSVA’^28^ through the single-sample gene set enrichment Analysis (ssGSEA) algorithm. Besides, the biological functions of each tumor sample were also quantified by GSVA, and the specific pathway signatures were derived and downloaded from the Hallmark gene sets in the MSigDB database (https://www.gsea-msigdb.org/gsea/msigdb/). Then, the ssGSEA pathway scores of cuproptosis together with several other gene features, including Apoptosis, Pyroptosis, Ferroptosis, TGF-β signal pathway, Interleukin-related pathway, Chemokine-related pathway, Chemokine Receptor-related pathway, Cell cycle pathway, Wnt signal pathway, mTOR signal pathway, Ras signal pathway, Akt signal pathway, EGF signal pathway, PI3K signal pathway, Metal Ion Homeostasis pathway, Fatty acid metabolism pathway, Glycolysis, and Glutathione metabolism were calculated by R package ‘GSVA’ through ssGSEA algorithm from MSigDB database^29^.

### Construction of Cuproptosis molecular phenotypes

Based on the expression level of 16 cuproptosis-related molecules, the unsupervised clustering analysis was used to identify the cuproptosis molecular phenotypes in colorectal cancer. The consensus clustering algorithm was applied to determine the optimal clustering numbers and phenotypes of CRC individuals and tumor cells in the TCGA-COREAD and the GSE132465 cohorts by the R package ‘ConsensusClusterPlus’ ^30^. Then, Principal Component Analysis (PCA) was used to show the cuproptosis-related molecule expression difference between different consensus phenotypes. In the single-cell levels, the differences in the expression distribution of the 16 cuproptosis-related molecules were analyzed and visualized by the R package “Seruat” ^31^. For the bulk RNA-seq data, the overall survival statistical difference between different phenotypes were calculated and visualized by R packages ‘survival’, ‘survminer’, and the variations in the expression levels of 16 cuproptosis-related molecules and the clinical characteristics between different phenotypes were shown by R package ‘ComplexHeatmap’ ^32^.

### Evaluation of TME immunological characteristics

Immunomodulators, including Major Histocompatibility Complex (MHC), immunoinhibitors, immunostimulators, and chemokines were collected from the previous study^33^. The anti-cancer immune system response has been proven with seven steps: (1) Release of cancer cell antigens, (2) Cancer antigen presentation, (3) Priming and activation, (4) Trafficking of immune cells to tumors, (5) Infiltration of immune cells into tumors, (6) Recognition of cancer cells by T cells, and (7) Killing of cancer cells. In this study, the gene data and the enrichment scores of each step in TCGA-COREAD cohorts were downloaded from the tracking tumor immune phenotype website (http://biocc.hrbmu.edu.cn/TIP/index.jsp) ^34–36^. Besides, the ssGSEA^33^, and CIBERSORT^37^ methods were used to estimate the immune cell infiltration of CRC individuals in the TCGA-COREAD cohorts and the Immune, Stromal, ESTIMATE scores, and tumor purity were calculated by R package ‘estimate’ ^38^. The HE staining immunophenotype pathology image data (Formalin-fixed paraffin-embedding) in TCGA datasets were obtained from the Cancer Digital Slide Archive (CDSA, https://cancer.digitalslidearchive.org/).

### Construction of the CuproScore

PCA algorithm was applied to construct a scoring system, called Cuproscore, to evaluate the levels of 16 cuproptosis-related molecules in CRC individuals. The positive and negative functions of 16 cuproptosis-related genes were calculated by univariate Cox regression according to the Cox coefficient. Then PCA algorithm was performed to analyze the expression levels of 16 cuproptosis-related molecules, and the principal component 1(PC1) was determined to present as the scores. The CuproScore formula is:

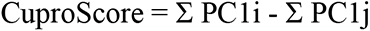

(Σ PC1i means the sum of PC1 of positive Cox coefficient cuproptosis molecules, and Σ PC1j means the sum of PC1 of negative Cox coefficient cuproptosis molecules). Then, the predictive values of CuproScore for the prognosis, molecular functions, and immunotherapy response of CRC individuals were further evaluated.

### Molecular and clinical significance of the CuproScore

Based on the median of CuproScore, the CRC patients in the TCGA-COREAD cohort were divided into the high and low CuproScore groups. Then the differences in the prognosis, mutation feature, molecular functions, and TME immunological characteristics between the two groups were also estimated by the same methods as previously introduced. The immunophenoscores, to predict the response to immunotherapy with CTLA-4 and PD-1 blockers in the TCGA datasets, were downloaded from the Cancer Immunome Database (TCIA) database(https://tcia.at/home) ^33^. To further explore the predictive value of CuproScore in anti-PD-L1, anti-PD1, and anti-CTLA4 immunotherapy, the CuproScore algorithm was also applied in the TCGA, IMvigor210, and Liu’s cohorts. And the individuals with stable disease (SD) or progressive disease (PD) as non-responders, while those with complete response (CR) or partial response (PR) as responders.

### High-throughput transcriptome analysis of CRC individuals

The study was approved by the Ethical Committee of Ruijin Hospital with a clinical trial registration number NCT04714814. Among the prospective patient cohort collected, 12 CRC individuals for the High-throughput transcriptome analysis were strictly screened in our center, without any medical treatment before recruitment. Detailed information about them is listed in Supplementary table S2. Total RNA was isolated from the surgically discarded normal and tumor colorectal tissues of these 12 CRC individuals. Then, the cDNA library was constructed, and high-throughput sequencing was performed at Shanghai Jiao Tong University School of Medicine (Shanghai, China) using the HiSeq 4000 (Illumina, San Diego, CA).

### Immunohistochemistry (IHC)

Cancer tissue-array including a total of 73-paired normal/tumor CRC individuals was also recruited from Ruijin Hospital (Shanghai, China). IHC training was performed according to standard protocols and the optimum sections of tissue specimens were obtained and deparaffinized. IHC was performed as the following antibodies: CDKN2A (RRID:AB_2861619), DLAT (RRID:AB_2761407), FDX1 (RRID:AB_2769449), and GCSH (RRID:AB_2760555). Each tissue-array was scored by three independent pathologists using a semi-quantitative method based on the German semi-quantitative scoring system^39^.

### Cell Lines and Cell Culture

NCM460 (RRID:CVCL_0460), HT29 (RRID:CVCL_0320), SW620 (RRID:CVCL_0547), HCT116 (RRID:CVCL_0291), SW480 (RRID:CVCL_0546), and RKO (RRID:CVCL_0504) cell lines were obtained from the American Type Culture Collection (ATCC, Manassas, VA). All cells were stored at the Shanghai Institute of Digestive Surgery and cultured in the corresponding medium with 10% fetal bovine serum (FBS), 0.1mg/ mL streptomycin, and 100U/ mL penicillin under the atmosphere at 37°C with 5% CO_2_.

### Cell viability assay

The cuproptosis inducer Elesclomol+Cu^2+^ (1:1 ratio, APExBIO) was dissolved in dimethyl sulfoxide (DMSO) to a total concentration of 40 mM. The working concentrations were diluted to 0, 0.75, 1.5, 3, 6, 12, 25, 50, 100, and 200 nM, and three wells were applied for each concentration. After being treated with gradient concentrations of Elesclomol+Cu^2+^, cell viability was measured by Cell Counting Kit 8 (CCK8, APExBIO), following the standard protocols. Then, the half-maximal inhibitory concentration (IC_50_) values of each cell line were determined using nonlinear regression.

### RNA Isolation and Real-Time PCR

RNA Isolation and Real-Time PCR were performed as previous methods^40^. The sequences of primers used for qPCR analysis were listed in Supplementary table S3. The relative mRNA expression levels of the 16 cuproptosis-related were calculated by the 2^−ΔΔCt^ method and normalized against that of *GAPDH*.

### Western bolt

Western blot was performed as previous methods^40^. Primary antibodies: β-Tubulin (RRID:AB_2861647, 1:2000), FDX1 (RRID:AB_2769449, 1:1000), and SLC31A1 (RRID:AB_2757632, 1:1000). Secondary antibody: anti-rabbit (RRID:AB_2819035, 1:5000), anti-mouse (RRID:AB_2877728, 1:5000).

### Statistical analysis

All the statistical analysis was conducted by the R software (version: 3.4.0) in this study. All P values of statistical data were based on two-sided statistical tests. P<0.05 was considered to be statistically significant. Spearman correlation coefficients were used to determine the correlation between variables. The unpaired student’s t-test was used to estimate the statistical significance of normally distributed variables, and the Mann-Whitney U test (also known as the Wilcoxon rank-sum test) was used to analyze non-normally distributed variables. To compare two or more groups, Kruskal-Wallis and one-way ANOVA tests were used. The "surv_cutpoint" function in the R package ‘Survminer’ was used to evaluate the critical value of each data set. The Kaplan-Meier method was used to generate survival curves for the subgroups of each data set, and the log-rank (Cox) test was used to determine statistically significant differences. A univariate Cox proportional hazard regression model was used to calculate the hazard ratio.

## Results

### Pan-cancer analysis of cuproptosis-related molecules (CPRMs)

The schematic flowchart of the study was presented in Figure 1. To explore the potential value of cuproptosis in tumors, the pan-cancer analysis of 16 cuproptosis-related molecules (CPRMs) was performed and the main results were as follows: Firstly, to evaluate the transcriptional changes between the normal and tumor tissues, the variations in the mRNA expression levels of the 16 CPRMs were shown in Figure 2A and all CPRMs were highly statistical significances in different tumor types in the TCGA pan-cancer cohorts, especially *CDKN2A*. Secondly, to estimate the genetic variation status of the 16 CPRMs in pan-cancer samples, a waterfall diagram shows the somatic mutation frequency and types of cuproptosis-related genes in Figure 2B. We found that *CDKN2A* had the highest mutation frequency of up to 4%, followed by *ATP7A* (2%) and *ATP7B* (2%) and the missense mutation was the most common type. The relationship between copy number variation (CNV) and mRNA expression levels of the 16 CPRMs in pan-cancer showed a positive correlation in most cancer types, especially in bladder urothelial carcinoma (TCGA-BLCA), breast invasive carcinoma (TCGA-BRCA), head and neck squamous cell carcinoma (TCGA-HNSC), Colon carcinoma (TCGA-COAD), Rectal carcinoma (TCGA-READ), and Lung squamous cell carcinoma (TCGA-LUSC) in Figure 2C. Then the prognostic values of cuproptosis-related genes in pan-cancer were also evaluated and the results showed that the CPRMs appeared to be the heterogeneous prognostic values in different cancers, such as most CPRMs played the protective roles in Kidney renal clear cell carcinoma (TCGA-KIRC) while mostly risky roles in brain lower-grade glioma (TCGA-LGG) in Figure 2D. Besides, in Figure 2E, we calculated the cuproptosis pathway score (ssGSEA) based on the 16 CPRMs and found there were also statistically significant differences in the score between the normal and tumor samples in pan-cancer, similarly to the situation of mRNA expression levels. Finally, to estimate the potential values of the cuproptosis pathway score, a systematic analysis was performed and a mostly negative correlation between the TME infiltrating cells and cuproptosis pathway score was found in pan-cancer except for the ovarian serous cystadenocarcinoma (TCGA-OV). In the molecular biology analysis, many pathways, such as adipogenesis, protein secretion, PI3K signaling, fatty acid metabolism, and oxidative phosphorylation were shown positively correlated with the cuproptosis pathway score in Figure 2F.

**Figure 1.**
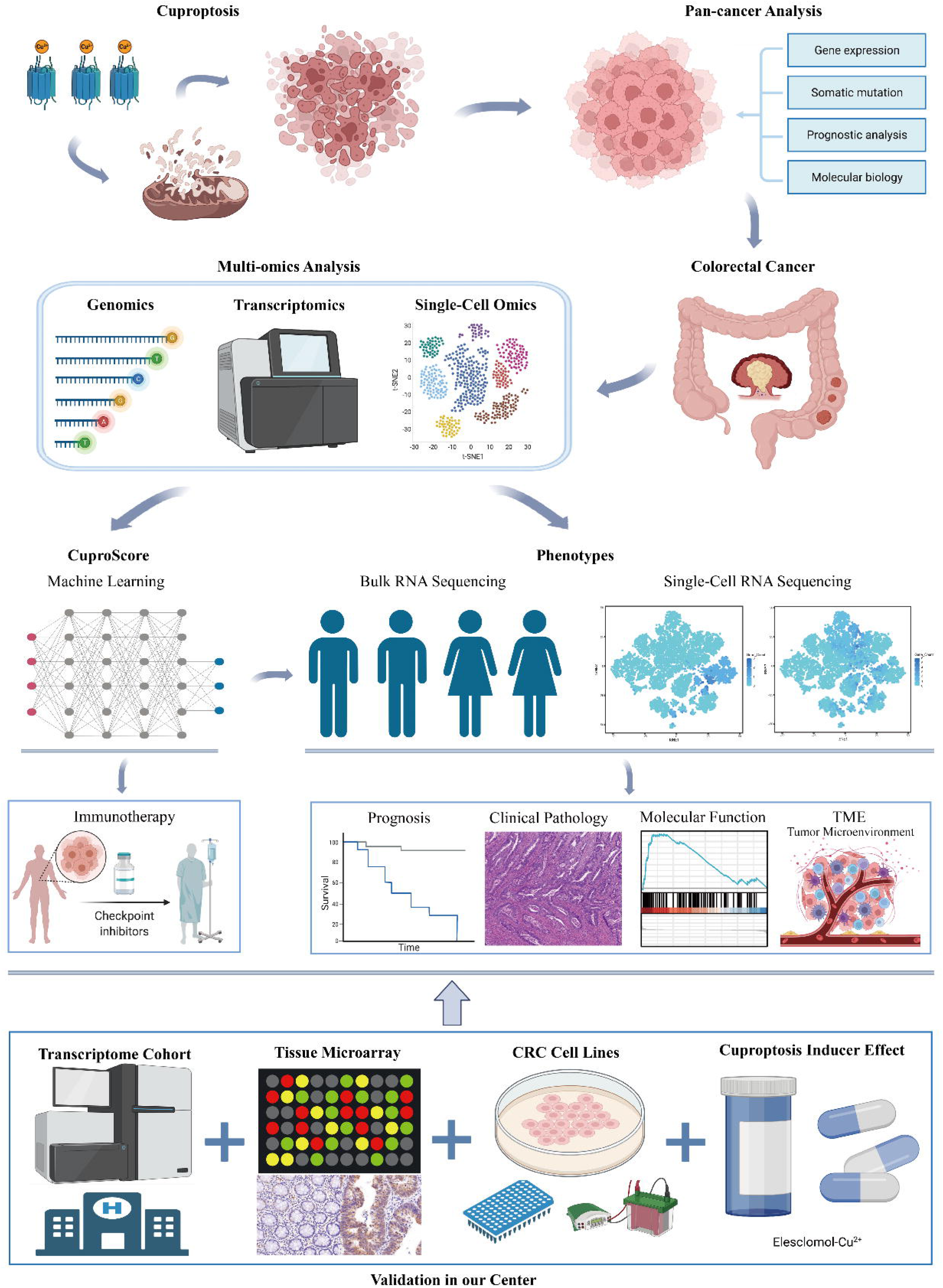
Schematic flowchart of the study design.

**Figure 2.**
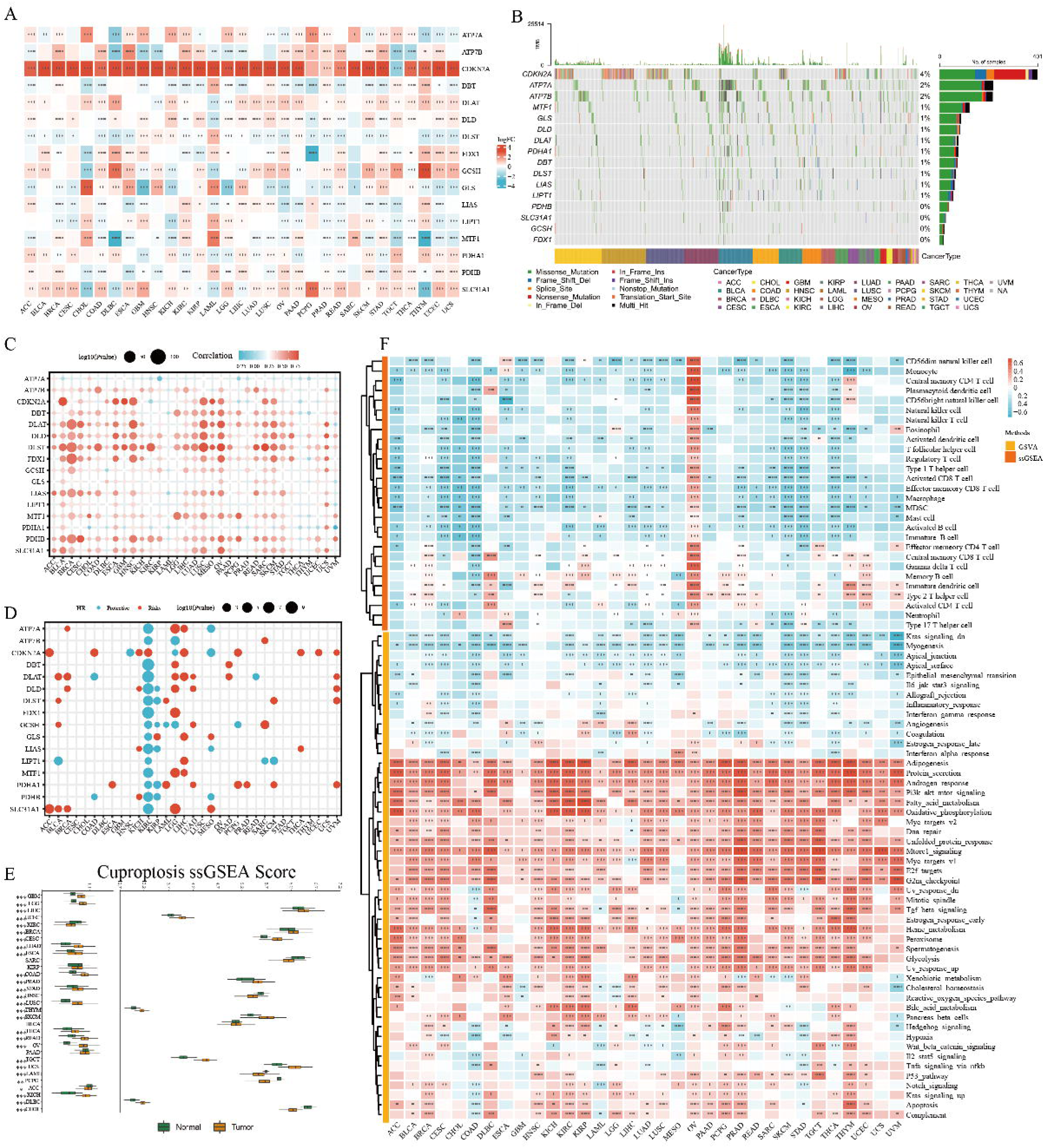
Pan-cancer analysis of cuproptosis-related molecules (CPRMs). A. The thermogram shows the different mRNA expression levels of 16 CPRMs between the normal and tumor samples in pan-cancer from TCGA cohorts. T-test: \**P* < 0.05, \*\**P* < 0.01, \*\*\**P* < 0.001. B. The waterfall diagram shows the somatic mutation frequency of 16 CPRMs in pan-cancer. C. The bubble chart shows the correlation between CNV and mRNA expression levels of 16 CPRMs in pan-cancer using Spearman correlation analysis. D. The bubble chart shows the relationship between prognosis and mRNA expression levels of 16 CPRMs in pan-cancer using the Log-rank test. E. The boxplot shows the different cuproptosis ssGSEA scores between the normal and tumor samples in pan-cancer. T-test: \**P* < 0.05, \*\**P* < 0.01, \*\*\**P* < 0.001. F. The thermogram shows the correlation between mRNA expression levels of 16 CPRMs and 28 TME infiltrating cells or 50 common molecular biology pathways in pan-cancer. Spearman correlation analysis: \**P* < 0.05, \*\**P* < 0.01, \*\*\**P* < 0.001.

### Clinical and Molecular Correlates of CPRMs in CRC

To further investigate the mRNA expression levels of 16 CPRMs in CRC bulk samples, a differential analysis was performed between the intestinal epithelium tissues and colorectal cancer tissues in GTEx and TCGA-COREAD cohorts. The results showed that each mRNA expression level of the 16 CPRMs was statistically different between normal and tumor tissues in Figure 3A, most of which were up-regulated in tumor samples, except *DBT* and *DLST*. In addition, we further validated the protein expression of the 16 CPRMs in HPA database (Supplementary Figure S1). Given that the cell lines may play an important role *in vitro* experiments, radar charts were applied to show the mRNA expression levels of 16 CPRMs in diverse CRC cell lines, demonstrating most CPRMs had sufficient expression abundance in CRC cell lines (Figure 3B). Then the somatic mutation frequency of CPRMs in 538 CRC individuals in the TCGA-COREAD cohorts was estimated in Figure 3C and found that *ATP7A* (4%) and *ATP7B* (3%) ranked top mutation frequency compared to others. However, the gain frequency of CNV in *ATP7A* was the least among CPRMs, while *ATP7B* ranked first in Figure 3D-E. To further explore the clinical values of CPRMs in CRC, the univariate Cox regression analysis indicated that *CDKN2A*, *DLAT*, *DLD*, and *PDHB* might have prognostic values in CRC individuals. *CDKN2A* tended to be a risk factor, and others contributed to the protective factors in Figure 3F. And the Kaplan-Meier curves showed the disease-specific survival (DSS), overall survival (OS), and progression-free survival (PFS) statistical discrepancy between high and low *CDKN2A* or *DLAT* expression groups in the TCGA-COREAD cohorts (Figure 3G). We also found that *LIAS*, *DLST*, *DLAT*, and *SLC31A1* had lower mRNA expression levels in the advanced TNM stage CRC compared to the early stage, while *ATP7B*, *CDKN2A*, and *GLS* had a higher level in stage IV CRC (Supplementary Figure S2A). Furthermore, based on the several drug resistance cohorts, we found the expression levels of many CPRMs were likely to be related to the chemotherapy, targeted therapy or immunotherapy resistance in CRC individuals, especially *DLAT* (Figure 3H). For the analysis of the molecular function, these CPRMs were mainly enriched in the oxidoreductase activity and metabolism-related pathways (Supplementary Figure S2B). For the molecular correlates, DLAT could be the hub molecule and had a positive correlation with other CPRMs (Supplementary Figure S2C-D). For the immune checkpoints analysis, there was a significant correlation between the CPRMs and immune checkpoint-related gene expression levels (Supplementary Figure S2E). For the ssGSEA and CIBERSORT analysis, the immune cell infiltration degree was negatively related to the expression levels of most CPRMs, except *DBT*, *DLST*, *MTF1* and *SLC31A1* (Figure 3I). For the GSVA analysis, *ATP7B*, *CDKN2A* and *PDHA1* had a negative correlation with most specific well-defined biological states or processes based on Hallmark gene sets, while other CPRMs were the opposite (Figure 3I).

**Figure 3.**
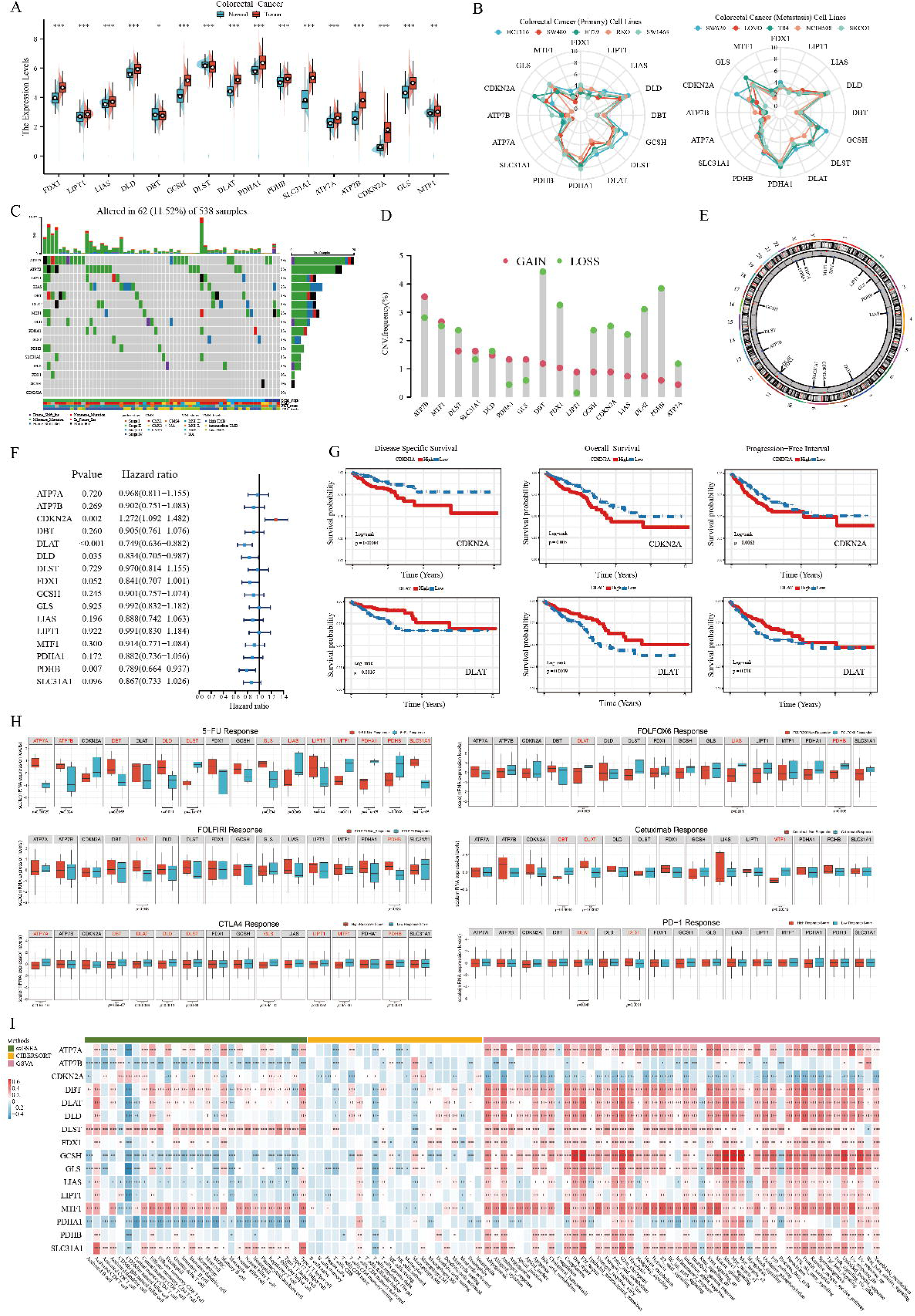
Expression and clinical significance of CPRMs in CRC. A. The boxplot shows the mRNA expression levels of 16 CPRMs between the normal and tumor CRC samples in GTEX and TCGA-COREAD cohorts. T-test: \**P* < 0.05, \*\**P* < 0.01, \*\*\**P* < 0.001. B. The radar charts show the mRNA expression levels of 16 CPRMs in different CRC cell lines in CCLE datasets. C. The waterfall diagram shows the mutation frequency of 16 CPRMs in 538 patients with colorectal cancer in the TCGA-COREAD cohorts. D. The vertical lollipop chart shows the frequencies of CNV gain, loss, and non-CNV among CPRMs in the TCGA-COREAD cohorts. E. The doughnut chart shows the locations of CNV alterations in CPRMs on 23 chromosomes. F. The forest map shows the results of Univariate regression analysis on the average survival rate of 16 CPRMs in the TCGA-COREAD cohorts. G. The Kaplan-Meier curves show disease-specific survival, overall survival, and progression-free interval of CDKN2A and DLAT in the TCGA-COREAD cohorts using the log-rank test. H. The boxplots show the relationship between mRNA expression levels of CPRMs and the chemotherapy, targeted therapy or immunotherapy resistance in CRC individuals. T-test: \**P* < 0.05, \*\**P* < 0.01, \*\*\**P* < 0.001. I. The thermogram shows the correlation between mRNA expression levels of 16 CPRMs and TME infiltrating cells or common molecular biology pathways in CRC individuals. Spearman correlation analysis: \**P* < 0.05, \*\**P* < 0.01, \*\*\**P* < 0.001.

### Identification of two cuproptosis-related molecular phenotypes (CMP) by unsupervised learning in CRC

To explore the role of cuproptosis in CRC individuals, we identified two cuproptosis-related molecular phenotypes (307 cases in CMP1 and 311 cases in CMP2) by unsupervised learning analysis using the R package ‘ConsensusClusterPlus’ (Figure 4A). The PCA analysis confirmed that the two phenotypes could be distinguished by the expression levels of the 16 CPRMs (Figure 4B). And patients in CMP1 had a worse prognosis than those in CMP2 in Figure 4C. The expression levels of cuproptosis molecules and the clinical characteristics were visualized using a thermogram in Figure 4D and there were significant differences in the expression levels of 15 CPRMs except DLST between the two phenotypes. To further understand the biological discrepancy between the two phenotypes, the GSVA analysis was performed and the results showed that the biological discrepancy mainly focused on the immune-, mechanism- and tumorigenesis-related pathways, such as Interleukin, PD-1, and CTLA4 signaling, Energy, Fatty acid, Glutathione, and Metal ions metabolism, and Cell cycle, p53, Wnt and Kras signaling in Figure 4E. In addition, there were also significant differences in the ssGSEA enrichment scores between the two phenotypes, including several Cell Death Pathways (Apoptosis, Ferroptosis, and Cuproptosis), Immune Pathways (TGF-β, Interleukin, and Chemokines), Tumorigenesis Pathways (Cell cycle, Wnt, mTOR, Ras, EGF, and PI3K), and Metabolism Pathways (Metal ion homeostasis, Fatty acid metabolism, and Glycolysis) in Figure 4F.

**Figure 4.**
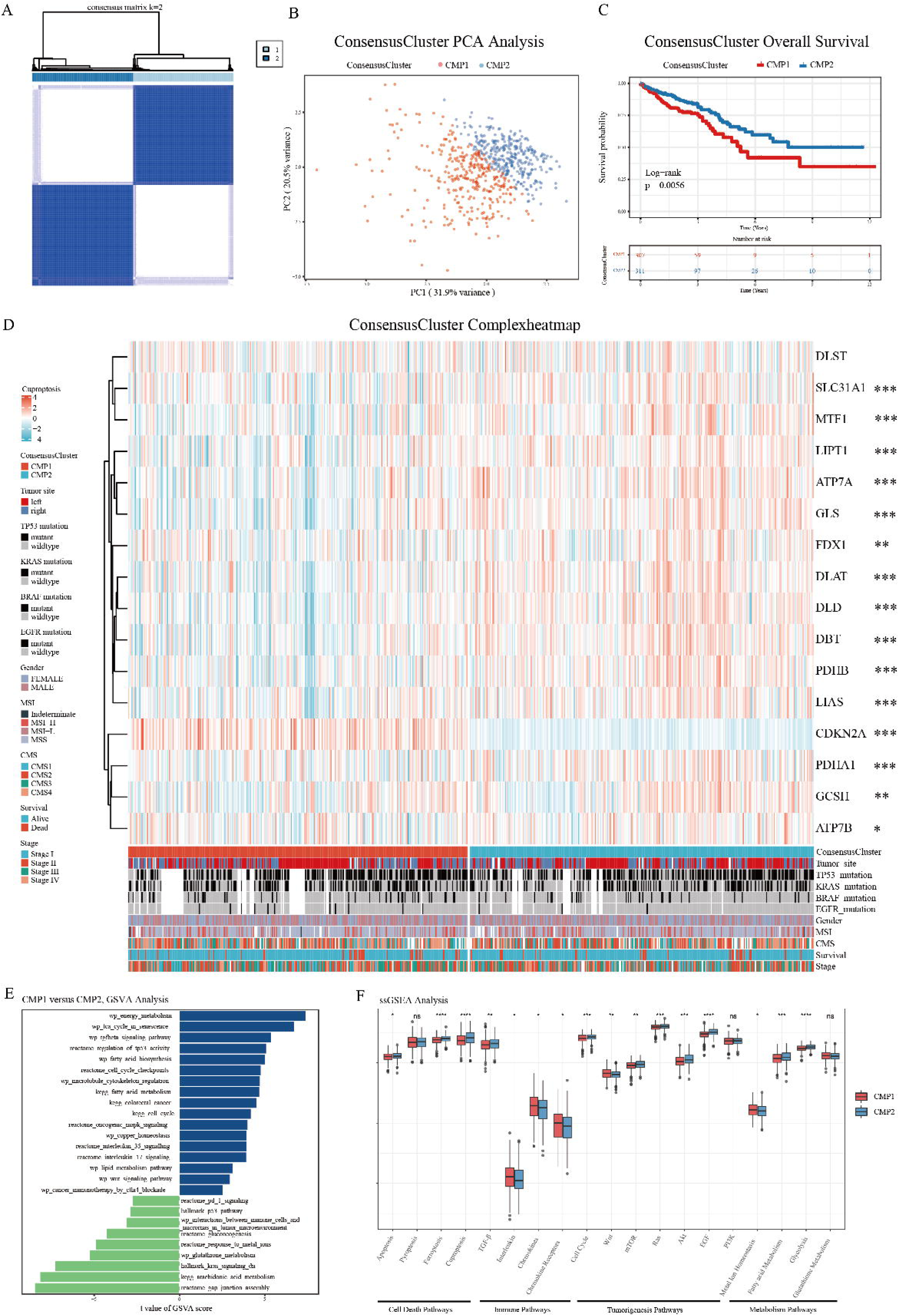
Identification of two cuproptosis-related molecular phenotypes by unsupervised learning in CRC. A. The consensus matrix thermogram defines two cuproptosis phenotypes (k = 2) and shows their correlation area by unsupervised learning. B. The principal component analysis of CPRMs shows a remarkable difference in transcriptomes between the two subtypes. C. The Kaplan-Meier curves for overall survival of two consensus phenotypes in TCGA using the log-rank test. D. The thermogram shows the differences in clinicopathologic features and mRNA expression levels of CPRMs between the two cuproptosis phenotypes. T-test: \**P* < 0.05, \*\**P* < 0.01, \*\*\**P* < 0.001. E. The bar chart shows the results of GSVA analysis of the differentially expressed genes between the two consensus phenotypes. F. The boxplot shows the differences in pathway ssGSEA scores between the two cuproptosis phenotypes. T-test: ^ns^*P* > 0.05, \**P* < 0.05, \*\**P* < 0.01, \*\*\**P* < 0.001.

### Variations in somatic mutation frequency and immune cell infiltration in two cuproptosis phenotypes

To explore the variations in somatic mutation frequency between two cuproptosis phenotypes, somatic mutation analysis was performed, and the results showed the different mutation frequencies of the common mutant genes in CRC between the CMP1 and CMP2, such as *APC* (72% vs 80%) and *KRAS* (45% vs 37%) in Figure 5A. Then the forest plot was drawn to show the different gene mutation distributions and samples in CMP1 were associated with higher mutation rates of several mutant genes, including *DKK4*, *CCDC15*, and *NTAN1*(Figure 5B). Besides, TMB levels were calculated and analyzed in both phenotypes, indicating that TMB levels were significantly higher in CMP1 subtype in Figure 5C. Based on the results of the above functional enrichment analysis, the immune process and pathways were significantly different between the two phenotypes. Thus, ESTIMATE, CIBERSORT, ssGSEA, and Cancer-immunity Cycle analysis were utilized to calculate the immune cell infiltration degree in each CRC sample based on the RNA-Sequencing data and explore the relationship between cuproptosis phenotype and TME. For the ESTIMATE analysis, samples in CMP1 had the higher immune and ESTIMATE scores which indicated that more immune components were contained (Figure 5D). For the CIBERSORT and ssGSEA analysis, it could be seen that samples in CMP1 had a higher infiltration degree of several immune cells with dominant anti-tumor activity, such as T cells, CD8^+^ T cells, central memory CD4^+^ T cell, T follicular helper cell, Type 17 T helper cell, NK cells, CD56dim natural killer cell, Monocyte, and Neutrophil (Figure 5E). Anti-tumor immune response has been proven as a series of step-by-step events referred to as the cancer-immune cycle. Our results indicated that these anti-tumor steps were mainly active in CMP1, including Step4 (Trafficking of immune cells to tumors): T cell recruiting, CD4 T cell recruiting, B cell recruiting, and Step5 (Infiltration of immune cells into tumors), while only Th22 cell recruiting in Step4 was enriched in CMP2 (Figure 4E). These results further confirmed the potential role of cuproptosis in tumor immune cell infiltration. Moreover, we found that CMP1 group had a higher expression of immunoinhibitors, including PD-1, LAG3, and TGFB1 (Figure 5F). MHC, immunostimulators, and chemokines were also more highly expressed in CMP1. In conclusion, our results further confirmed that more infiltration of immune cells was contained in the tumor tissue of CMP1 group, while less in that of CMP2 group.

**Figure 5.**
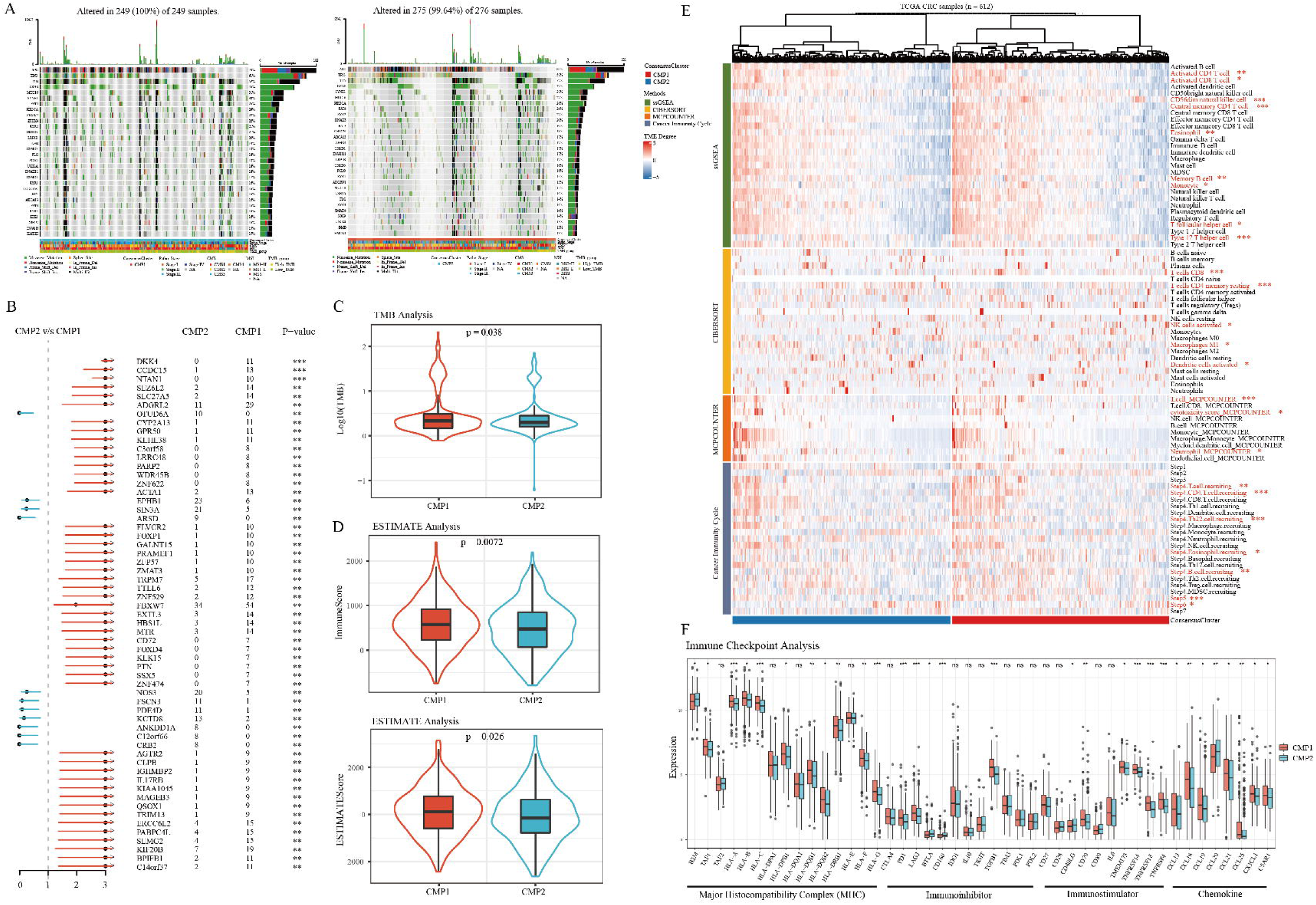
Variations in somatic mutation frequency and immune cell infiltration in two cuproptosis phenotypes. A. The waterfall diagrams show genetic alterations of common mutant genes between the two cuproptosis phenotypes. B. Forest plot shows variations in the somatic mutation frequencies between the two cuproptosis phenotypes. Chi-square test: \*\**P* < 0.01, \*\*\**P* < 0.001. C. The violin plot shows variations in TMB scores between the two cuproptosis phenotypes. T-test: \**P* < 0.05. D. The violin plots show variations in TME scores between the two cuproptosis phenotypes using ESTIMATE analysis. T-test: \**P* < 0.05, \*\**P* < 0.01. E. The thermogram shows variations in immune scores and the frequency of TME infiltrating cells between the two cuproptosis phenotypes. Wilcoxon test: \**P* < 0.05, \*\**P* < 0.01, \*\*\**P* < 0.001. F. The boxplot shows variations in mRNA expression of different kinds of immune checkpoints between the two cuproptosis phenotypes. Wilcoxon test: ^ns^*P* > 0.05, \**P* < 0.05, \*\**P* < 0.01, \*\*\**P* < 0.001.

### Single-cell transcriptome analysis of CRC cells based on cuproptosis-related molecular phenotypes

Considering the influence of different cell populations in bulk tissue, the differential analysis of the single-cell RNA cohort was performed to investigate the single-cell mRNA expression levels of 16 CPRMs in CRC cells. Consistent with the bulk-seq results, significant expression differences were demonstrated between 1070 normal and 17469 tumor epithelial cells, as shown in Figure 6A. Besides, based on the results of the differential analysis, the t-SNE plots were presented to show the molecular expression of these CPRMs at the single-cell level in Figure 6C. Then, to explore whether the cuproptosis-related molecular classification could be also exactly reflected in the CRC single cells, 17,469 CRC epithelial cells were screened from the single-cell cohort, and then also identified two cuproptosis-related molecular phenotypes (7408 single cells in CMP1-like and 10061 single cells in CMP2-like) by unsupervised learning analysis using the R package ‘ConsensusClusterPlus’ in Figure 6B. To verify the variations in the proportion and number of molecular and clinical characteristics between the two cuproptosis-related single-cell molecular phenotypes, the stacked histograms were shown in Figure 6D. For clinical significance, tumor cells in CMP1-like were predominantly derived from TNM stage III&IV, right-hemi colon cancer, poorly or mucinous differentiation CRC individuals. For the CMS subtype, tumor cells in CMP1-like were mainly derived from CMS1, CMS3, or CMS4 subtype CRC individuals through both single-cell and bulk levels. In contrast, tumor cells in CMP2-like were enriched in the CMS2 subtype which indicated a better prognosis. For MSI subtypes, tumor cells in CMP1-like were predominantly clustered into the MSI-H subtype, which intimated a higher probability of sensitivity to immunotherapy. For gene mutation type, tumor cells in CMP1-like were mainly clustered into the KRAS mutation group while tumor cells in CMP2-like were predominantly clustered into APC and TP53 mutation groups. Consistent with the functional enrichment analysis in bulk levels, we found the biological discrepancy was also focused on the immune-, mechanism- and tumorigenesis-related pathways between the two cuproptosis-related single-cell molecular phenotypes, such as ATP metabolic process, immune response, oxidative stress, IL-17 signaling, Glutathione metabolism, Glycolysis and Cell cycle pathways by GO and KEGG analysis in Figure 6E-F.

**Figure 6.**
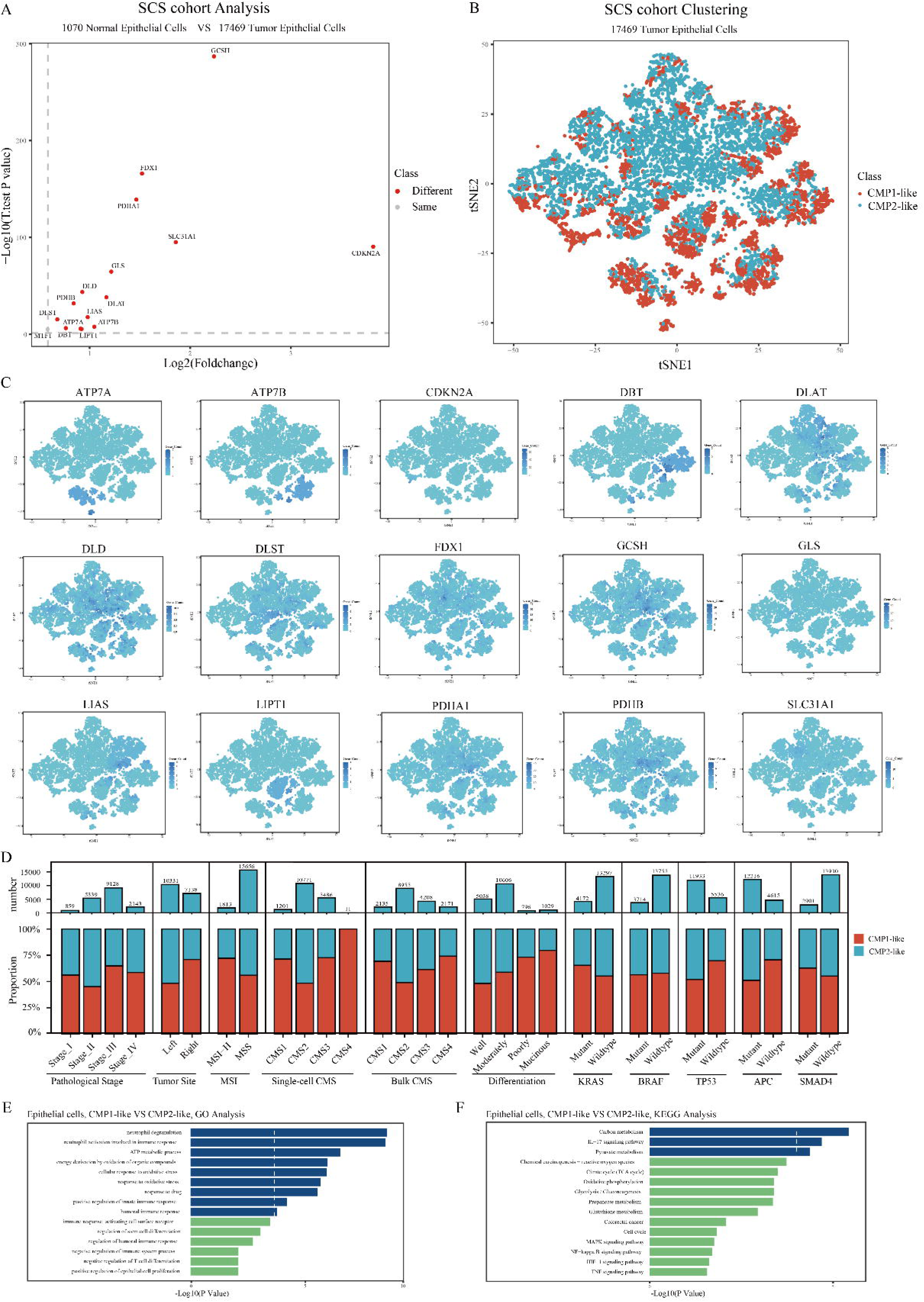
Single-cell transcriptome analysis of CRC cells based on cuproptosis-related molecular classification. A. The volcano map shows the different single-cell mRNA expression levels of 16 CPRMs between 1070 normal and 17469 tumor epithelial cells. B. The t-SNE visualization shows the different single-cell distributions between the two cuproptosis phenotypes. C. The t-SNE visualizations show the molecular expression of 16 CPRMs except MTF1. D. The stacked histogram shows variations in the proportion and number of molecular and clinical characteristics between the two cuproptosis single-cell phenotypes. E-F. The bar plots show the results of GO (E) and KEGG (F) analysis of the differentially expressed genes between the two cuproptosis single-cell phenotypes.

### Construction and evaluation of a novel cuproptosis-related scoring system (CuproScore) in CRC

As the above results, cuproptosis may play an important role in the regulation of tumorigenesis, TME, and survival in colorectal cancer. Thus, based on the 16 CPRMs, a novel cuproptosis-related scoring system called CuproScore was constructed to quantify cuproptosis regulation in each individual using the PCA algorithm. In the TCGA-COREAD cohorts, samples in CMP1 had a higer CuproScore than those in CMP2 in Figure 7A. Meanwhile, the high CuproScore group, divided by median or optimal cut-off score, all had the worse prognosis and the Kaplan-Meier curves were shown in Figure 7B-C. For the somatic mutations, different mutation frequencies of the common mutant genes were presented between high and low CuproScore groups, such as APC (72% vs 81%) and KRAS (44% vs 38%), as shown in Figure 7D. Besides, we also found that the CuproScore of the high TMB group was higher than that of the low TMB group in Figure 7E. For the molecular pathways, the GSVA analysis was further performed and the results showed that the biological discrepancy also mainly focused on the immune-, mechanism- and tumorigenesis-related pathways, such as Interleukin, TGF-β, and CTLA4 signaling; Energy, Fatty acid, copper homeostasis, and lipid metabolism; Cell cycle, p53, mTOR, Wnt, and Kras signaling in Figure 7F. For the TME, we evaluated the correlation between CuproScore and immune cell infiltration degree and found that CD56dim natural killer cell, monocyte, and central memory CD4 T cell had a statistically positive correlation with cuproptosis-related scores, while Th22 cell recruiting tended to be negative in Figure 7G. In addition, for the ESTIMATE analysis, the CuproScore also had a statistically positive relationship with TMEscore (Immunescore and ESTIMATEscore) in Figure 7H. The HE staining immunophenotype also confirmed that the high CuproScore group displayed a higher lymphocyte infiltration and further suggested that there was a positive correlation between cuproptosis and immune cell infiltration in Figure 7I.

**Figure 7.**
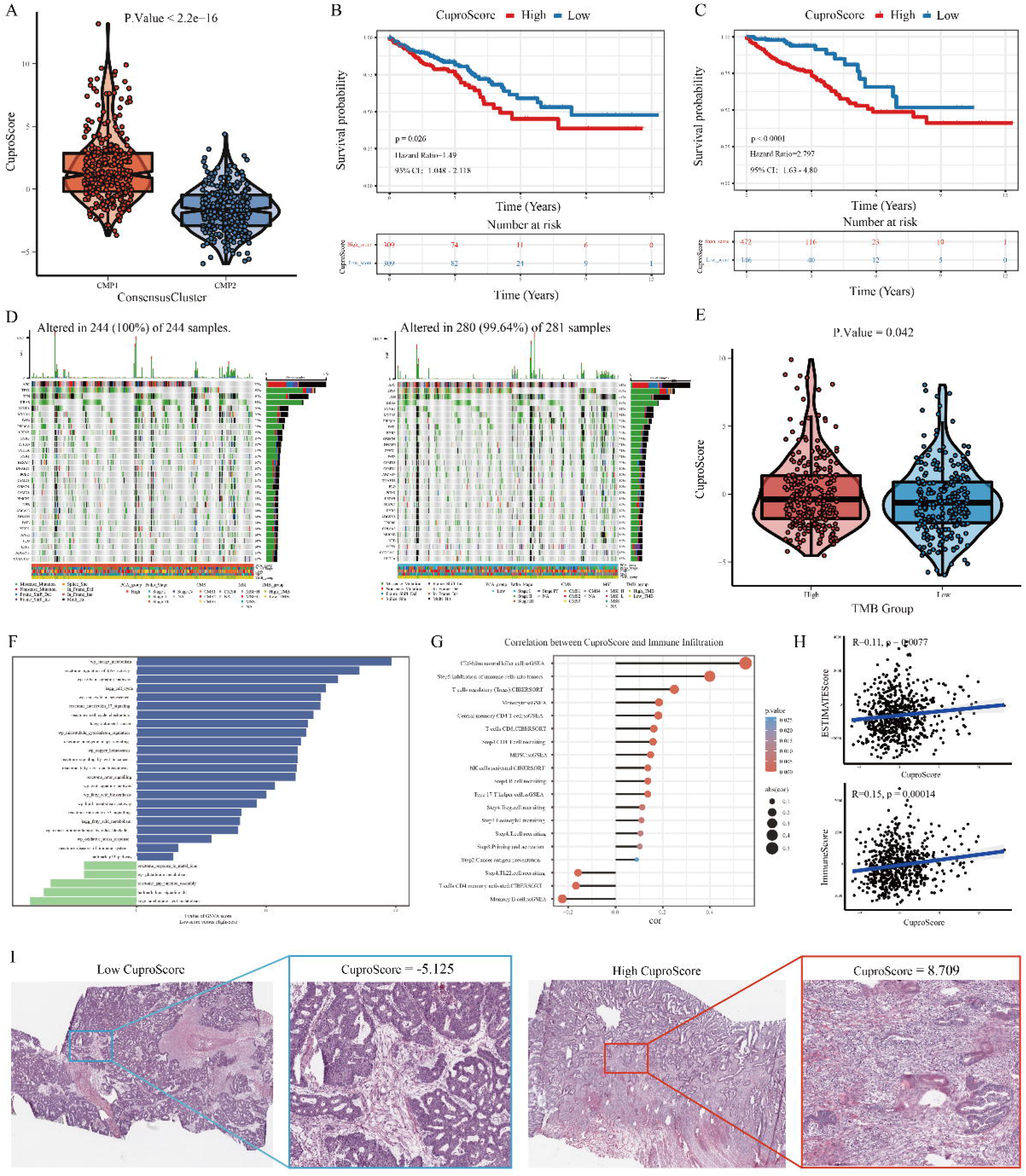
Construction and evaluation of a novel cuproptosis-related scoring system (CuproScore) in CRC. A. The violin plot shows variations in CuproScore between the two cuproptosis phenotypes using the Wilcoxon test. B-C. The Kaplan-Meier curves show the significant difference in the survival rate between the high and low CuproScore groups in the TCGA database using median (B) or best (C) cut-off values. D. The waterfall diagrams show genetic alterations of common mutant genes between the high and low CuproScore groups. E. The violin plot shows variations in CuproScore between the high and low TMB groups using the Wilcoxon test. F. The bar chart shows the results of GSVA analysis of the differentially expressed genes between the high and low CuproScore groups. G. The lollipop chart shows the correlation between the CuproScore and the level of immune cell infiltration. H. The scatter plots show the correlation between the CuproScore and TME scores (ESTIMATE and Immune scores) using Spearman correlation analysis. I. The representative images show the variations in pathological HE staining between the high and low CuproScore groups.

### Predictive value of CuproScore for immunotherapy response

Immunotherapies represented by anti-PD-L1, anti-PD1, and anti-CTLA4 have been widely applied in clinical treatments. Considering the molecular and clinical significance of CuproScore in CRC as described above, especially in immune cell infiltration, we next selected several immunotherapy-related cohorts to investigate whether the CuproScore could predict an effective response to these immunotherapies in different types of cancer. For the TCGA-COREAD cohorts, the results showed significant variations in CuproScore between different immunotherapy groups with high or low predicted immunotherapy sensitivity scores in Figure 8A. Then, for the IMvigor210 cohort, patients with high CuproScore had a significantly better prognosis compared to those with low CuproScore (Figure 8B). In addition, patients with high CuproScore were more likely to get an effective clinical response to anti-PD-L1 immunotherapy (Figure 8C-D). Furthermore, the CuproScore of the immune-inflamed phenotype was significantly higher than that of the immune-desert and immune-excluded phenotypes (Figure 8E). And the result indicated the CuproScore could be negatively correlated with the rejection of treatment and the desert immunophenotype. It has been proven that the expression level of PD-L1 was strongly correlated with the efficacy of anti-PD-L1 immunotherapy. Thus, the relationship between the CuproScore and the tumor-infiltrating immune cells (IC) and tumor cells (TC) immune types was further analyzed in Figure 8F-G and found the CuproScore was positively correlated with PD-L1 expression in tumor cells. For Liu’s cohort, consistent with the previous conclusions, the CuproScore could also predict the prognosis of melanoma individuals and their clinical response to the anti-PD-1 immunotherapy (Figure 8H-I). Taken together, the CuproScore we constructed could be a potentially powerful scoring system for predicting prognosis and the response to immunotherapies in different cancers.

**Figure 8.**
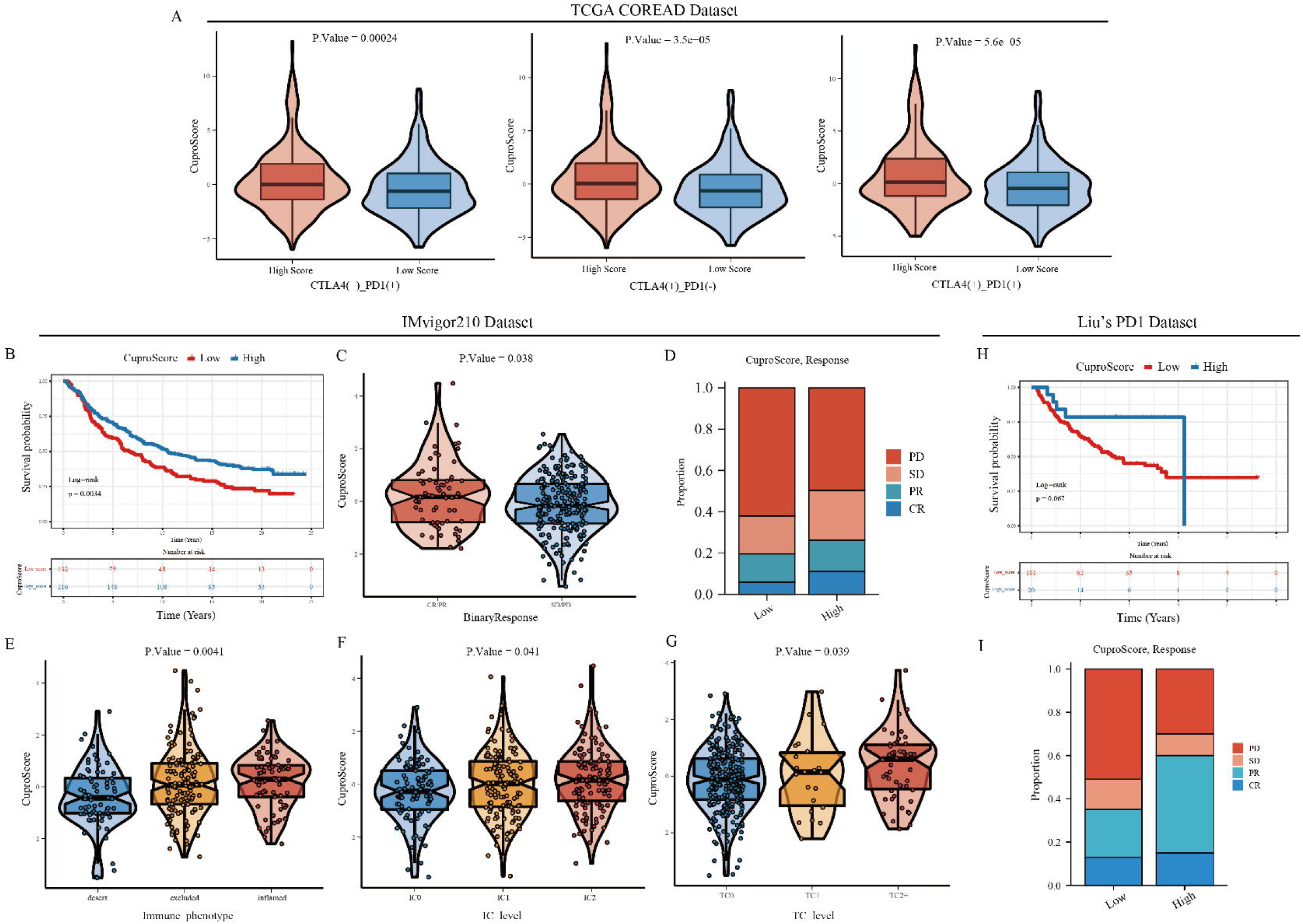
Predictive value of CuproScore for immunotherapy response. A. The violin plots show variations in CuproScore between groups with high or low immunotherapy sensitivity scores in the TCGA-COREAD cohorts. B. The Kaplan-Meier curve shows the significant difference in the survival rate between the high and low CuproScore groups in the IMvigor210 cohort using the best cut-off value. C. The violin plot shows variation in CuproScore between groups with different anti-PD-L1 responsiveness. D. The stacked histogram shows variations in the proportion of different anti-PD-L1 responsiveness between the high and low CuproScore groups. E. The violin plot shows variations in CuproScore among three immune phenotypes. F-G. The violin plots show variations in CuproScore among PD-L1 expression of different IC (F) and TC (G). H-I. The Kaplan-Meier curve (H) and stacked histogram (I) show the potential predictive value of CuproScore for anti-PD-L1 immunotherapy in Liu’s dataset.

### Validation of the clinical and molecular value of CuproScore in our transcriptome cohort

Considering over-reliance on the public datasets could destabilize the results, we constructed our transcriptome cohort (12 CRC individuals from Ruijin hospital) to further verify the value of CuproScore. The CuproScore of each sample was calculated based on the above scoring system, and the median CuproScore was utilized to classify samples into high or low CuproScore groups. The clinical characteristics of all these samples were presented in Supplementary Table S2. Consistent with the above results in the TCGA-COREAD cohorts, CRC individuals in the high CuproScore group tended to have higher immune cell infiltrations, including active CD4 T cells, active CD8 T cells, and CD56bright natural killer cells (Figure 9A). Meanwhile, the immune checkpoint expression levels, such as PD-1, PD-L1, and CTLA4, were also higher in the high CuproScore group (Figure 9B), and CuproScore had a statistically positive correlation with the immune cell infiltrations (Figure 9C) and immune checkpoint expression (Figure 9D). For the GO and KEGG analysis, the differentially expressed genes between the high and low CuproScore groups were similarly enriched in the immune-, mechanism- and tumorigenesis-related pathways, such as regulation of immune response, T/B cell receptor signaling pathway, and CD8 positive alpha beta t cell differentiation; Fatty acid, Copper homeostasis, and Inositol phosphate metabolism; Cell cycle, p53, mTOR, and VEGF signaling pathways (Figure 9E-F). For the GSEA analysis, Wnt-β-catenin signaling and Hypoxia were suppressed in the high CuproScore group while several inflammation-related functions, including interferon γ response and IL6 JAK STAT3 signaling pathways, were up-regulated (Figure 9G). The HE staining immunophenotype also confirmed that the high CuproScore group displayed a higher lymphocyte infiltration (Figure 9H).

**Figure 9.**
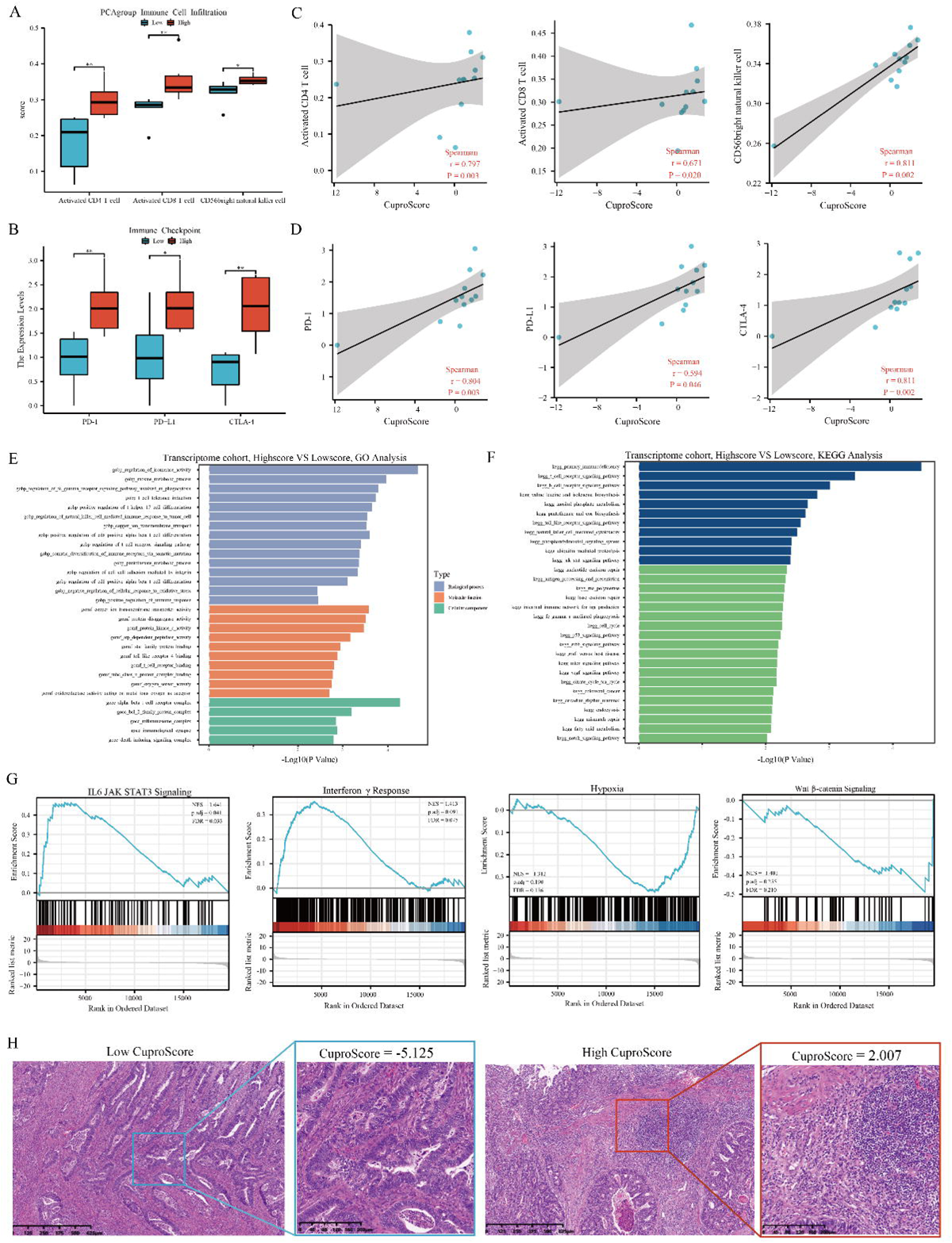
Validation of the clinical and molecular value of CuproScore in our transcriptome cohort. A-B. The boxplot shows the differences in immune cell infiltration (A) and immune checkpoint expression levels (B) between low and high CuproScore groups in CRC individuals from our transcriptome cohort (Ruijin hospital). Wilcoxon test: \**P* < 0.05, \*\**P* < 0.01, \*\*\**P* < 0.001. C-D. The scatter charts with regression lines show the correlation between CuproScore and immune cell infiltration (C), and immune checkpoint expression levels (D) using Spearman correlation analysis. E-F. The bar charts show the results of GO (E) and KEGG (F) enrichment analysis between the high and low CuproScore groups by GSVA. G. The string diagrams show the enrichment pathways between low and high CuproScore groups using GSEA analysis based on the differential expression genes. H. The representative images show the variations in pathological HE staining between the high and low CuproScore groups from our transcriptome cohort.

### Validation of the expression and clinical significance of 16 CPRMs in our cohorts

To address the scientificity and reliability of the above results, the transcriptome cohort, cell lines, and tissue-array of our center (Ruijin hospital) were applied to verify the expression levels and clinical significance of 16 CPRMs in CRC. For the transcriptome cohort, the differential analysis was performed between the paired normal and tumor tissues from 12 CRC individuals in our center. The results showed that the mRNA expression levels of *FDX1, LIPT1, DLD, GCSH, DLAT, PDHA1, PDHB, SLC31A1, ATP7A, ATP7B, CDKN2A, GLS,* and *MTF1* were statistically different between normal and tumor tissues in Figure 10A and all of them were up-regulated in tumor samples consistent with the above results in the TCGA-COREAD cohorts. Then, for the cell line level, the normal colon cell line (NCM460) and five CRC cell lines (HT29, SW620, HCT116, SW480, and RKO) were utilized to verify the 16 CPRMs expression levels in Figure 10B. Different from the results in TCGA and Ruijin cohorts, the expression levels of *DBT* and *DLST* were significantly up-regulated, while *ATP7B* was downregulated in CRC cell lines, compared to normal colon cell lines. Furthermore, the protein expression levels and clinical significance of hub CPRMs, including FDX1, DLAT, CDKN2A, and GLS were further verified in the tissue-arrays of our center by IHC (Figure 10C-E). Consistently, the results showed that the expression of FDX1, DLAT, CDKN2A, and GLS all increased in tumor tissues. Meanwhile, DLAT and CDKN2A were also associated with the prognosis of CRC individuals in our center.

**Figure 10.**
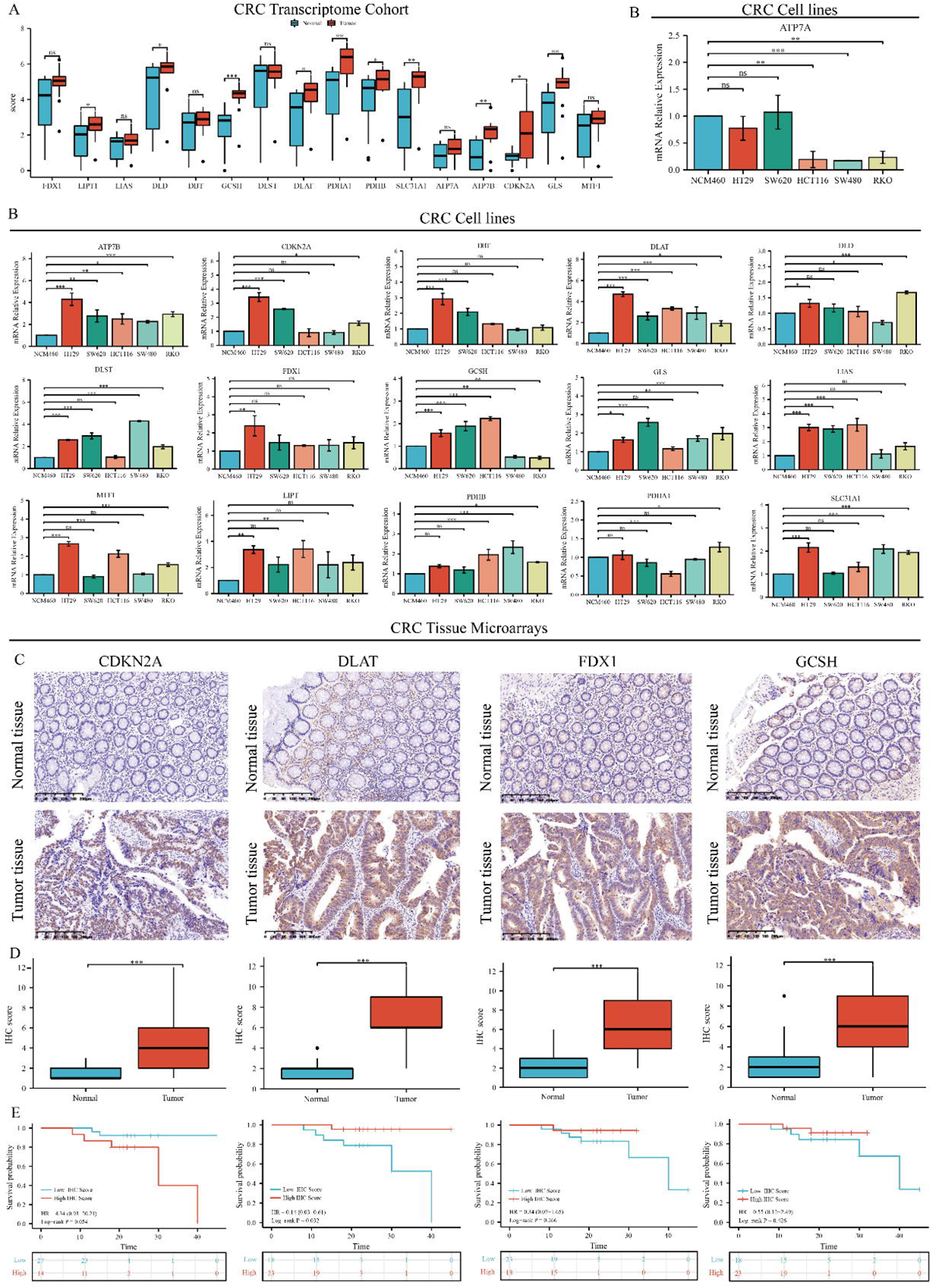
Validation of the expression and clinical significance of 16 CPRMs in our cohorts. A. The boxplot shows the different mRNA expression levels of 16 CPRMs between the normal and tumor CRC samples from our transcriptome cohort. T-test: \**P* < 0.05, \*\**P* < 0.01, \*\*\**P* < 0.001. B. The boxplots show the different mRNA expression levels of 16 CPRMs between the normal colon cell line (NCM460) and five CRC cell lines (HT29, SW620, HCT116, SW480, and RKO) using real-time PCR. One-way ANOVA: ^ns^*P* > 0.05, \**P* < 0.05, \*\**P* < 0.01, \*\*\**P* < 0.001. C. Representative immunohistochemistry images show the different protein expression levels of CDKN2A, DLAT, FDX1, and GCSH in CRC tissues and corresponding normal tissues from the CRC tissue-array of Ruijin hospital. D. The boxplots display the differences in IHC scores for CDKN2A, DLAT, FDX1, and GCSH between CRC tissues and corresponding normal tissues from the CRC tissue-array. Wilcoxon test: \*\*\**P* < 0.001. E. The Kaplan-Meier curves show overall survival of CDKN2A, DLAT, FDX1, and GCSH in the CRC tissue-array using the log-rank test.

### The effects of cuproptosis inducer Elesclomol-Cu^2+^ on 16 CPRMs in CRC cell lines

To explore the role of 16 CPRMs in cuproptosis, CRC cell lines HT29 and SW620 were treated with different working concentrations of the cuproptosis inducer Elesclomol-Cu^2+^. The copper-chelating agent D-penicillamine was acted as the cuproptosis inhibitor and used to perform the restoration experiments of cuproptosis. The results of the cell viability assay indicated that Elesclomol-Cu^2+^ could effectively inhibit the cell proliferation of HT29 and SW620 underlying a dose-dependent manner. The IC_50_ of each cell line was all-around 30 nM (Supplementary Figure S3). Based on their IC_50_, different concentrations of Elesclomol-Cu^2+^ (15, 30, and 45 nM) were applied to treat HT29 and SW620 cell lines for 24h. In addition, after 30 nM concentration of Elesclomol-Cu^2+^ treatment for 24 h, 30 µM concentration of D-penicillamine was then used to treat these cell lines for 12h. Then, the mRNA expression levels of 16 CPRMs in these cell lines were investigated by Real-Time PCR. The results indicated that the mRNA expression levels of 11 CPRMs (*FDX1, LIPT1, LIAS, DLD, DBT, GCSH, DLAT, PDHA1, PDHB, ATP7A,* and *ATP7B*) were decreased and 5 CPRMs (*DLST, SLC31A1, CDKN2A, GLS,* and *MTF1*) were increased after being treated with Elesclomol-Cu^2+^ in Figure 11A. However, the mRNA expression level of *DLD* was not statistically different in the SW620 cell line. After the D-penicillamine treatment, the variations in the mRNA expression levels of most CPRMs in both two cell lines were reversed and seemed to be returned to normal levels, while a few of them had no significant statistical difference. In addition, consistent with the mRNA expression results, the different protein expression levels of the hub CPRMs (FDX1 and SLC31A1) were further confirmed by Western Blot and shown in Figure 11B. In summary, the potential effects of cuproptosis inducer Elesclomol-Cu^2+^ on 16 CPRMs in CRC could be verified by *in-vitro* experiments.

**Figure 11.**
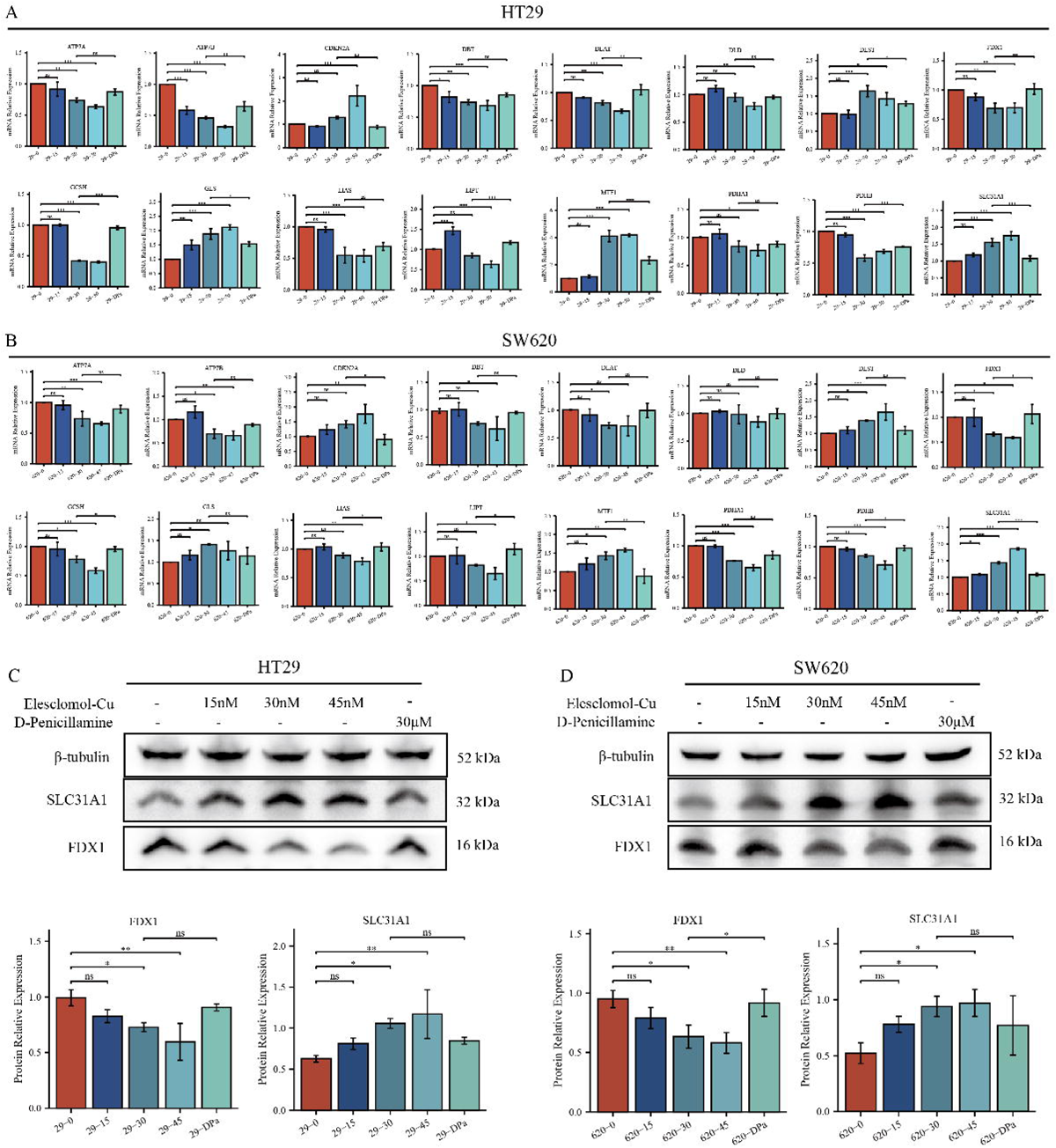
The effects of cuproptosis inducer Elesclomol-Cu^2+^ on 16 CPRMs in CRC cell lines. A-B. The boxplots show the different mRNA expression levels of 16 CPRMs in HT29 (A) and SW620 (B) cell lines after being treated with 15, 30, 45nM Elesclomol-Cu^2+^ and 30μM D-penicillamine using real-time PCR. C-D. The western blots and boxplots show the different protein expression levels of hub CPRMs (FDX1 and SLC31A1) in HT29 (C) and SW620 (D) cell lines after being treated with 15, 30, 45nM Elesclomol-Cu^2+^ and 30μM D-penicillamine using Western Blot. One-way ANOVA: ^ns^*P* > 0.05, \**P* < 0.05, \*\**P* < 0.01, \*\*\**P* < 0.001.

## Discussion

Recently, besides apoptosis, new forms of regulated cell death have been identified, including necroptosis^41^, pyroptosis^42^, Autophagy^43^, etc. Among them, heavy metal-related RCD has drawn more and more attention in this field, such as ferroptosis and cuproptosis^10, 44^. Heavy metals are essential for normal function, however, like a double-edged sword, either too little or too much can lead to cell death. Tresv demonstrated that cuproptosis, copper-induced RCD, was featured by aggregation of lipoylated mitochondrial enzymes and the destabilization of Fe–S cluster proteins^10^. However, how copper specifically induces cell death and the potential value of cuproptosis and cuproptosis-related molecules in clinical treatments remain largely unknown.

In the current study, we aimed to comprehensively explore the role of cuproptosis in CRC. Pan-cancer analysis was applied to explore the molecular and clinical significance of 16 CPRMs in 33 cancer types, including transcriptional changes, genetic variation status, prognostic differences, and molecular functions. Then, TCGA (TCGA-COREAD), GEO (GSE62080, GSE19862, GSE19860, GSE108277, and GSE81005), and single-cell RNA sequencing (GSE132465) and immunotherapy-related (IMvigor210 and Liu’s) cohorts were integrated to systematically evaluate the association between cuproptosis and tumorigenesis, TME, and immunotherapy response in CRC. Two CMPs were identified in both bulk and single-cell RNA sequencing samples in CRC, which were closely related to clinicopathological and TME cell-infiltrating characteristics. Furthermore, a scoring system called CuproScore was established to quantify the expression rate of CPRMs in the individual tumor sample. We further confirmed these results using transcriptome cohort of CRC tissues, cell lines, and tissue-array of our center.

Copper is an essential microelement to maintain normal physiological^45^. At the same time, the copper imbalance is closely associated with a variety of human diseases, including Wilson’s disease^46^, cardiovascular disorders^47^, and cancers^48^. Due to the unique energy and metabolic demands in tumors, copper has been proven to play dual roles in tumor initiation and tumor progression, especially in TME. Recently, Tsvetkov et al. have found a novel form of cell death caused by copper ionophores, called cuproptosis. Sixteen CPRMs have been identified to regulate the process of cuproptosis. Indeed, heavy metal-related RCD is not uncommon. Ferroptosis, an iron-dependent cell death, was firstly reported in 2012^9^ and a growing number of studies have evidenced the potential value of ferroptosis and ferroptosis-related molecules in tumors, especially in tumorigenesis, tumor therapy, and tumor immunity^49–51^. Distinct from ferroptosis, cuproptosis is another type of ion-metabolism-related RCD, which was dependent on mitochondrial respiration and the TCA cycle, rather than ROS or mitochondrial ROS^52^. In the current study, the pan-cancer analysis showed that the CPRMs expression was deregulated between normal and tumor tissues in 33 cancer species, which also closely associated with both clinical prognosis and molecular functions, consistent with the previous study^53^.

Copper is mainly (50–70%) absorbed by the colonic epithelium^54^. Compared to the normal colonic epithelium, the copper metabolism has been reprogrammed and the expression levels of copper transport and metabolism molecules are also distinctly changed in CRC, leading to the severe imbalance of intratumoral copper and facilitating tumor progression, invasion, and metastasis^55^. Copper overload has been proven to be a profound inducer of cuproptosis^10^. Therefore, it could be more meaningful to explore the relationship between cuproptosis and CRC. In this study, most CPRMs were found up-regulated in CRC, and four of them (CDKN2A, DLAT, DLD, and PDHB) were demonstrated with prognostic value. Meanwhile, the expression levels of LIAS, DLST, DLAT, SLC31A1, and ATP7B were related to the TNM stages in CRC individuals. Furthermore, CPRMs expression levels were associated with the pathological characteristics, tumorigenesis-related pathways, tumor immune microenvironment, and tumor resistance in CRC. Surprisingly, we found that DLAT could be a potential marker in CRC, especially in drug-resistance. CRC individuals with higher DLAT expression are more likely resistant to chemotherapy and immunotherapy. DLAT, a subunit of the mitochondrial pyruvate dehydrogenase complex^56^, plays an important role in tumorigenesis and carbohydrate metabolism, acting as a core metabolic node that promotes the TCA cycle and mediates pyruvate oxidation in cancers^57^. The aggregation of DLAT may also lead to the cuproptosis in lung cancer cells^10^. In our study, the potential value of DLAT has been demonstrated in CRC individuals and it could be meaningful to further explore its specific mechanism in the future.

As a result of the integration of molecular changes, molecular classification has been emerging as a useful tool in clinical treatments for tumor patients. The status of MSI-H/dMMR or CMS of CRC patients is highly significant for therapy option and prognosis prediction^58^. Based on the robust advancement of Next Generation Sequencing technology and machine learning, by which we constructed the cuproptosis-related molecular subtyping in this study, CMP1, and CMP2. We claimed that there were obviously differences in molecular functions, TNM stages, and prognosis between these two phenotypes of CRC patients. The differentially expressed genes between two phenotypes were obviously enriched in the tumor immune-related and tumor metabolism-related pathways, indicating diverse TME existed in different subtypes. The results of single-cell transcriptome analysis also confirmed a similar tendency. Despite the recent progress in tumor immunotherapy, there is a great heterogeneity in the efficacy of immunotherapy in CRC individuals, which highlights that CRC may have a special TME different from other tumors. TME is a complex tissue structure composed of the extracellular matrix, blood vessels, immune cells, and fibroblasts^59^. Immune cells, the major cellular components of TME, include various types of anti-tumor immune cells (T cells, B cells, NK cells, and Dendritic cells) and pro-tumor immune cells (Tregs, MDSC, and Macrophages) ^60, 61^, which have been proven to play a significant role in tumor development, progression and therapeutic resistance. Therefore, it is meaningful to further explore the potential influence of cuproptosis on TME. TMB is an important indicator to evaluate the effect of immunotherapy and is positively correlated with immunotherapy response to tumor individuals^62^. In our results, we found that CRC individuals in CMP1 were presented with higher TMB than those in CMP2.

Increasing evidence has shown that the accumulative infiltration of T cells, NK cells, and Dendritic cells suggests a positive role in the immune defense of CRC^63, 64^. Besides, a recent study revealed that B cells also play an important role in therapeutic responses to ICIs by altering T cell activation and function^65^. We observed more infiltration of immune cells (CD4^+^ T cell, CD8^+^ T cell, NK cell, B cell, Dendritic cell, and Neutrophil et.al.) in CMP1, indicating cuproptosis was involved in the tumor immune microenvironment and has a stronger immunoactivated effect of CRC individuals in CMP1. ICIs are a revolutionary breakthrough in tumor therapy, usually, higher expression of PD-1 or PD-L1 indicates better sensitivity to ICIs. We also found individuals in CMP1 have higher expression levels of ICIs, indicating better immunotherapy response. Unfortunately, the relationship between tumor immune microenvironment and the prognosis of tumor patients is still controversial, especially in CRC^66, 67^. In our study, we found that CRC individuals in CMP1 had a poor prognosis while higher immune cell infiltrations. To our mind, the possible reasons for this result are as follows: (1) CRC has a unique tumor and immune microenvironment different from other solid tumors, leading to the small proportion of CRC individuals benefiting from immunotherapy^68, 69^; (2) The immune microenvironment among tumor patients is highly variable, which is even heterogenous in different regions of the same tumor tissue^70^. Evidence showed that nearly one-third of tumors show different immune infiltration^71^; (3) Anti-tumor activity is dependent on tumor antigen presentation and tumor cell recognition^72^. It is necessary to clarify whether the infiltrated immune cells can recognize antigens and play a role in killing tumors. (4) With the detailed IHC analysis of solid tumors, three classifications of tumor immunophenotypes have emerged: ‘hot’ or ‘immune-inflamed’ tumors (with pronounced immune cell infiltrations in the tumor core), ‘immune-excluded’ tumors (showing immune infiltrates at the tumor boundaries), and ‘cold’ or immune-desert’ tumors (with no immune cell infiltration in tumor) ^73, 74^. While recent studies have confirmed that the average immune cell infiltration around the tumor is higher than in the tumor core^75, 76^, the concept of tumor immunophenotypes has been ignored in these studies. Meanwhile, the common transcriptome/bulk sequencing technique cannot effectively distinguish ‘immune-inflamed’ and ‘immune-excluded’ tumors. Several spatially resolved ways or techniques, such as spatial transcriptomics, will be performed to close the knowledge gap in the future.

To quantify the expression level of 16 CPRMs in each patient as much as possible, a scoring system, CuproScore, was also constructed based on the PCA algorithm, which was further comprehensively utilized in several tumors, including CRC, bladder and kidney cancer, to assess its clinical value. Our results demonstrated patients in CMP1 often presented higher CuproScore, consistent with the molecular phenotype of cuproptosis (CMP1 and CMP2), higher CuproScore was closely related to worse prognosis, higher TMB and more immune cell infiltration. Similar to the results in the TCIA dataset, higher CuproScore indicated better sensitivity to anti-PD-1, anti-PD-L1, or anti-CTLA-4 ICIs in the TCGA dataset. Unfortunately, there was no research cohort of immunotherapy available with open access in CRC. Thus, we checked the predictive capability of CuproScore in other tumor immunotherapy. The same trends of CuproScore were also observed in IMvigor210 and Liu’s cohorts. Therefore, we speculate that cuproptosis may be related to immunotherapy in different types of tumor patients and the combination of cuproptosis inducers and ICIs could be a potential therapeutic strategy. In summary, CuproScore could be applied to evaluate the infiltration characteristics of TME cells, predict the prognosis of individuals, and distinguish which patients may benefit from immunotherapy in different types of tumors, especially in CRC.

Despite the recent developments in bioinformatics that have provided great convenience in biological science research, there are still limitations by this method, such as over-reliance on the public databases, lack of enough biological and clinical validation, and ignoring differences in ethnicities. Therefore, to ensure our study is more scientific and convincing, we used the tissue transcriptome, the tissue-array, and the *in vitro* experiments of our cohort to verify the above results. According to our transcriptome cohort and HE staining immunophenotype results, CRC individuals in the high CuproScore group may have more immune cell infiltrations (CD4^+^ T cell, CD8^+^ T cell, and NK cell), higher expressions levels of immune checkpoint (PD-1, PD-L1, and CTLA-4) and more lymphocyte infiltrations of pathological specimens. Based on the results of enrichment analysis, the differentially expressed genes were similarly enriched in the immune-, mechanism- and tumorigenesis-related pathways. We also used the tissue-array and the *in vitro* experiments to verify the expression and clinical significance of 16 CPRMs in our center. It is noteworthy that DLAT and CDKN2A were associated with the prognosis of CRC individuals in both public and our cohorts. Considering the little understanding of the role of cuproptosis in cancers, whether cuproptosis inducers could be developed as effective anti-tumor therapies, such as Elesclomol-Cu^2+^, etc., remains largely unknown. Therefore, we applied Elesclomol-Cu^2+^ to trigger cuproptosis in two CRC cell lines, HT-29 and SW-620. And we detected significant lethal toxicity in both CRC cell lines by Elesclomol-Cu^2+^ at low concentration, revealing IC_50_ value was both around 30 nM. To evaluate whether these 16 CPRMs play key roles in cuproptosis of CRC, the expression levels of CPRMs were further tested after treatment with different concentrations of Elesclomol-Cu^2+^. Not surprisingly, most of these CPRMs were dysregulated at mRNA or protein levels, while the underlying mechanism remained unknown. Therefore, cuproptosis inducer could be a novel therapeutic strategy in CRC and these CPRMs could be potential targets for inducing tumor cell cuproptosis personalized anti-tumor therapy. As mentioned above, mitochondrial respiration was required for cuproptosis, while glucose-induced glycolysis showed less sensitivity to cuproptosis. Therefore, there is a hint to future tumor therapy that inhibition of glucose-related metabolism may sensitize tumor cells to cuproptosis while still impairing the tumor’s malignant behavior. In addition, the Elesclomol, or other copper ionophores, may be of benefit to the tumors with high expression of lipoylated mitochondrial proteins, or with tolerance to conventional apoptosis-induced therapy. And we expect that our study could promote the development of new therapeutic or combined therapeutic strategies in the future.

Several limitations have been recognized in our study. The novel CMP and scoring system (CuproScore) were constructed based on the retrospective data from several databases, and further prospective studies are necessary to testify their clinical value. To minimize the influence of tumor heterogeneity on outcomes, we have used different databases, including public datasets, our transcriptome cohort, and tissue-array, there could be still some impact. Therefore, we are also collecting an expanded cohort of patients from multiple centers to validate these results. Although the tumor killing effects of Elesclomol-Cu^2+^ has been proven in our study, the effectiveness of the combination of cuproptosis inducer with other drugs, such as chemotherapy drugs, targeted drugs or ICIs, for CRC patients is still unclear and meaningful to be further clarified.

In summary, our study suggested that cuproptosis played an important role in CRC development, remodeling tumor microenvironment and response to immunotherapy. Cuproptosis could be a promising therapeutic target and a useful tool for therapeutic interventions and possible combination treatments in CRC patients.

## Availability of Data and Materials

All the public datasets can be downloaded in the Genomic Data Commons (GDC, https://portal.gdc.cancer.gov/), the University of California Santa Cruz (UCSC, https://xenabrowser.net/datapages/), the Cancer Cell Line Encyclopedia project (CCLE, https://portals.broadinstitute.org/ccle), the Human Protein Atlas (HPA, https://www.proteinatlas.org/) database, and the Gene Expression Omnibus (GEO, https://www.ncbi.nlm.nih.gov/geo/) database. RNA-Seq Data will be deposited in NCBI SRA after all projects are completed but it can be available from the authors upon reasonable request.

## Acknowledgements

We appreciate the patients who have participated in TCGA, GEO, and Ruijin cohorts.

## Author Contributions

S.Z., M.Z., and J.S. served as corresponding authors and designed the study. Y.S. and X.F. contributed equally to this work and should be considered co-first authors. X.Y., S.L., X.Z., and L.H. charged the data curation and coordinated the research activities. Y.S. and X.F. collected the data, performed the major analysis and interpreted the data. S.Z. and J.S. supervised the study. S.Z., M.Z., and J.S. verified the replication and reproducibility of results. Z.S., Y.S., and X.F. drafted the manuscript. All authors have read and agreed to the published version of the manuscript.

## Additional Files

### Supplementary Figures

Supplementary Figure S1: The protein expression of the 16 CPRMs in HPA database in CRC.

Supplementary Figure S2: Clinical values and interaction of Cuproptosis-related molecules (CPRMs) in CRC. A. The boxplots show the expression differences of 16 CPRMs between different clinical stages in CRC. B. The bar charts show the enrichment pathways of 16 CPRMs from KEGG and GO databases. C. The sector graph indicate the correlation of 16 CPRMs by mRNA expression levels. D. The Protein-Protein Interaction network plot show the interaction of 16 CPRMs from STRING database. E. The heatmap indicate the correlation between the expression levels of 16 CPRMs and immune-related genes, including MHCs, immunostimulators, immunoinhibitors.

Supplementary Figure S3: The IC50 of HT29 and SW620 CRC cell line with Elesclomol-Cu^2+^ treatment.

Supplementary Figure S4: Compound structures for Elesclomol and D-Penicillamine in the study.

Supplementary Figure S5: Source Western-blot Images for Figure 11C.

Supplementary Figure S6: Source Western-blot Images for Figure 11D.

### Supplementary Tables

Supplementary Table S1: The full name, function of the 16 CPRMs.

Supplementary Table S2: The clinical characteristics of the individuals in our transcriptome cohort.

Supplementary Table S3: Real-time PCR primer sequences of 16 CPRMs.

