## Supplementary Data for "Cuproptosis is correlated with clinical status, tumor immune microenvironment and immunotherapy in colorectal cancer: a multi-omic analysis"

#### Supplementary Figures

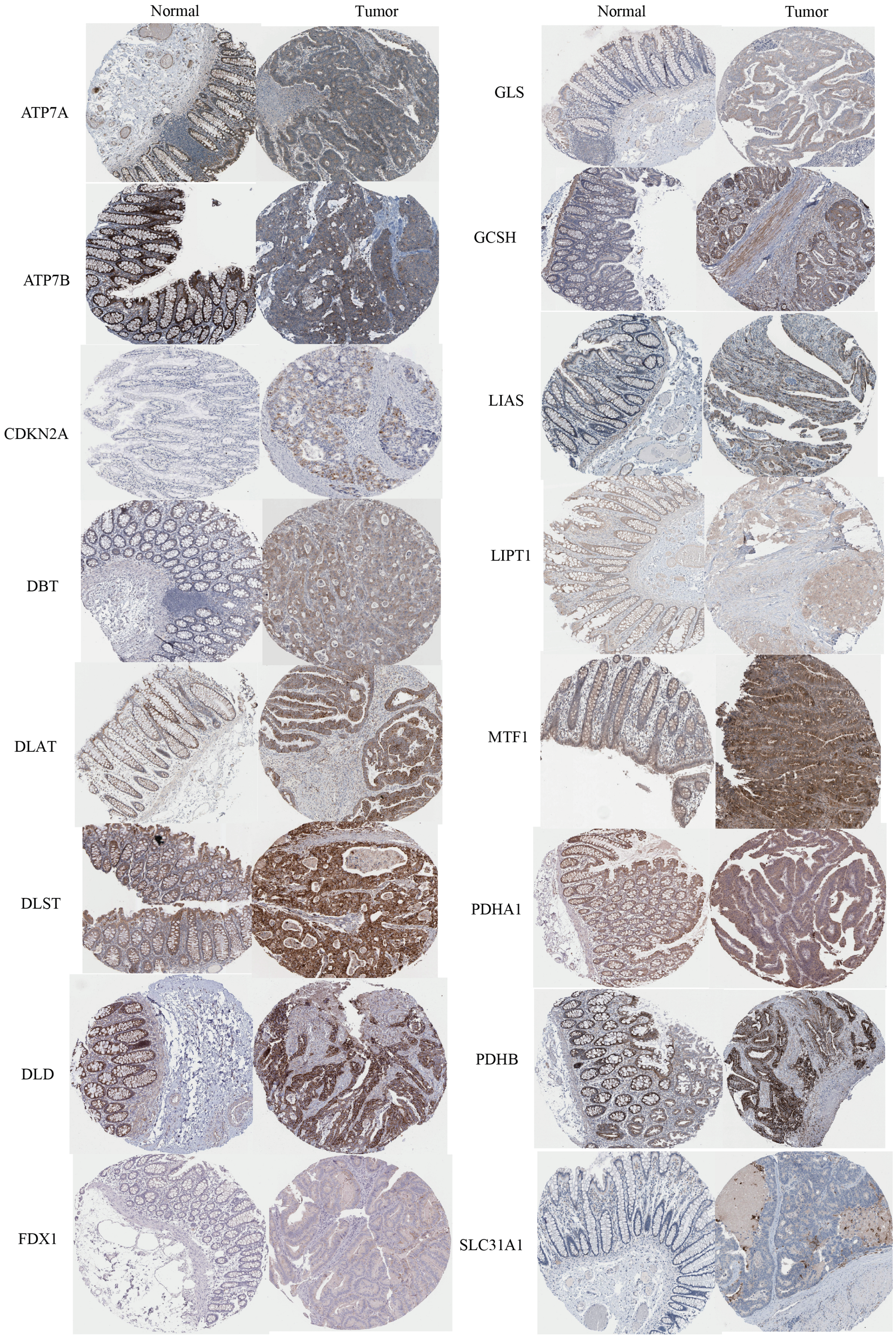


**Figure S1 The protein expression of the 16 CPRMs in HPA database in CRC**

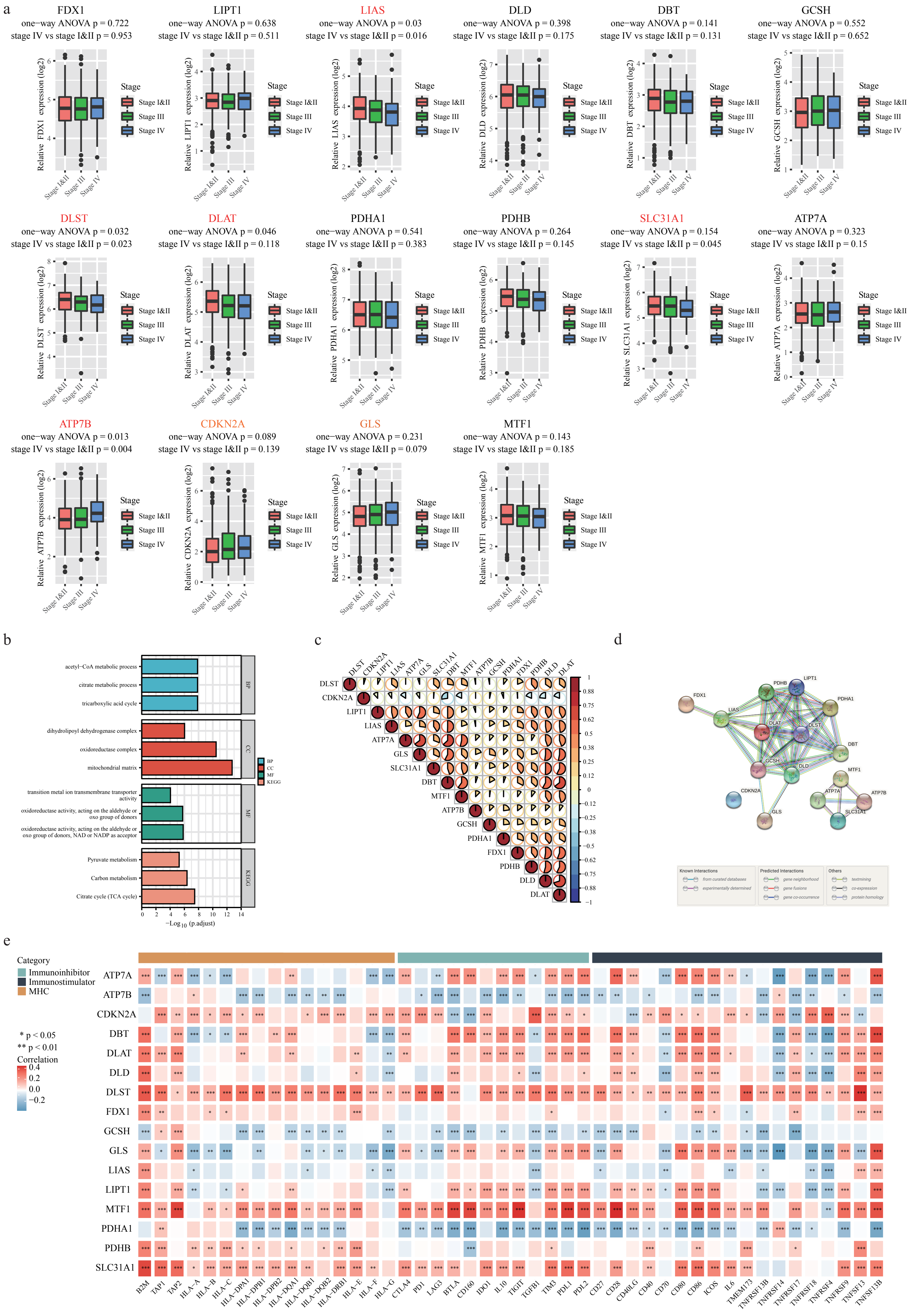
a
FDX1
one−way ANOVA p = 0.722
stage IV vs stage I&II p = 0.953
LIPT1
one−way ANOVA p = 0.638
stage IV vs stage I&II p = 0.511
LIAS
one−way ANOVA p = 0.03
stage IV vs stage I&II p = 0.016
DLD
one−way ANOVA p = 0.398
stage IV vs stage I&II p = 0.175
DBT
one−way ANOVA p = 0.141
stage IV vs stage I&II p = 0.131
GCSH
one−way ANOVA p = 0.552
stage IV vs stage I&II p = 0.652
DLST
one−way ANOVA p = 0.032
stage IV vs stage I&II p = 0.023
DLAT
one−way ANOVA p = 0.046
stage IV vs stage I&II p = 0.118
PDHA1
one−way ANOVA p = 0.541
stage IV vs stage I&II p = 0.383
PDHB
one−way ANOVA p = 0.264
stage IV vs stage I&II p = 0.145
SLC31A1
one−way ANOVA p = 0.154
stage IV vs stage I&II p = 0.045
ATP7A
one−way ANOVA p = 0.323
stage IV vs stage I&II p = 0.15
ATP7B
one−way ANOVA p = 0.013
stage IV vs stage I&II p = 0.004
CDKN2A
one−way ANOVA p = 0.089
stage IV vs stage I&II p = 0.139
GLS
one−way ANOVA p = 0.231
stage IV vs stage I&II p = 0.079
MTF1
one−way ANOVA p = 0.143
stage IV vs stage I&II p = 0.185
Stage
Stage I&II
Stage III
Stage IV
b
acetyl−CoA metabolic process
citrate metabolic process
tricarboxylic acid cycle
dihydrolipoyl dehydrogenase complex
oxidoreductase complex
mitochondrial matrix
transition metal ion transmembrane transporter activity
oxidoreductase activity, acting on the aldehyde or oxo group of donors
oxidoreductase activity, acting on the aldehyde or oxo group of donors, NAD or NADP as acceptor
Pyruvate metabolism
Carbon metabolism
Citrate cycle (TCA cycle)
BP
CC
MF
KEGG
−Log10 (p.adjust)
c
d
Known Interactions
Predicted Interactions
Others
e
Category
Immunoinhibitor
Immunostimulator
MHC
* p < 0.05
** p < 0.01
Correlation

**Figure S2 Clinical values and interaction of Cuproptosis-related molecules (CPRMs) in CRC**
a.The boxplots show the expression differences of 16 CPRMs between different clinical stages in CRC.b. The bar charts show the enrichment pathways of 16 CPRMs from KEGG and GO databases. c. The sector graph indicate the correlation of 16 CPRMs by mRNA expression levels. d. The Protein-Protein Interaction network plot show the interaction of 16 CPRMs from STRING database. e. The heatmap indicate the correlation between the expression levels of 16 CPRMs and immune-related genes, including MHCs, immunostimulators, immunoinhibitors**.**

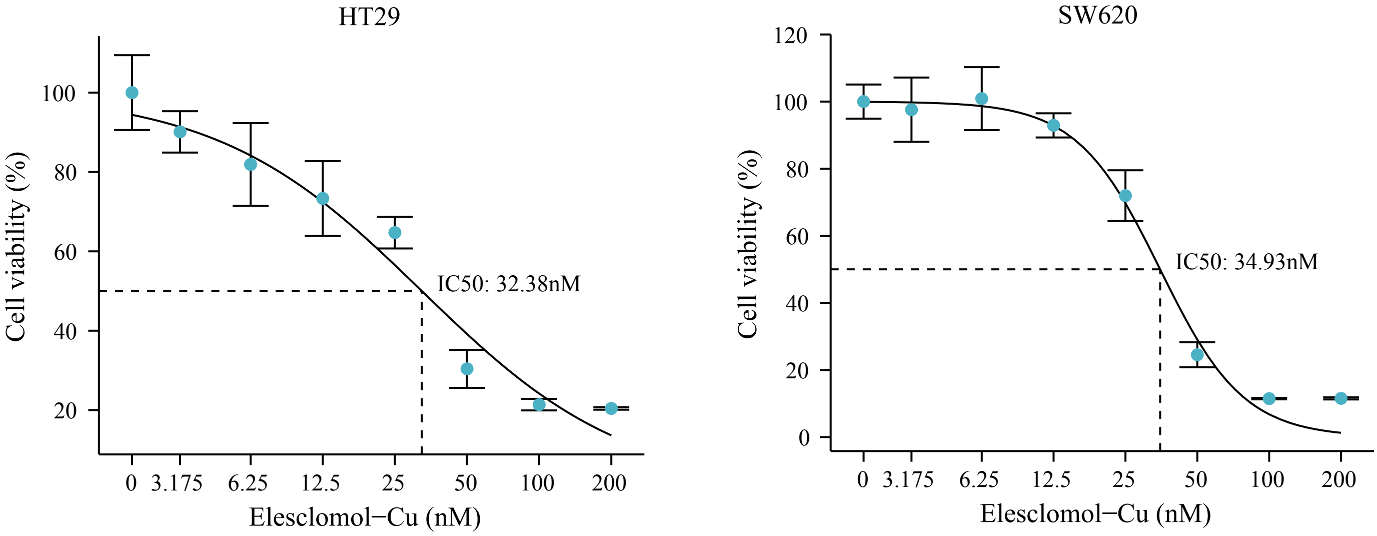


**Figure S3 The IC50 of HT29 and SW620 CRC cell line with Elesclomol-Cu2+ treatment**

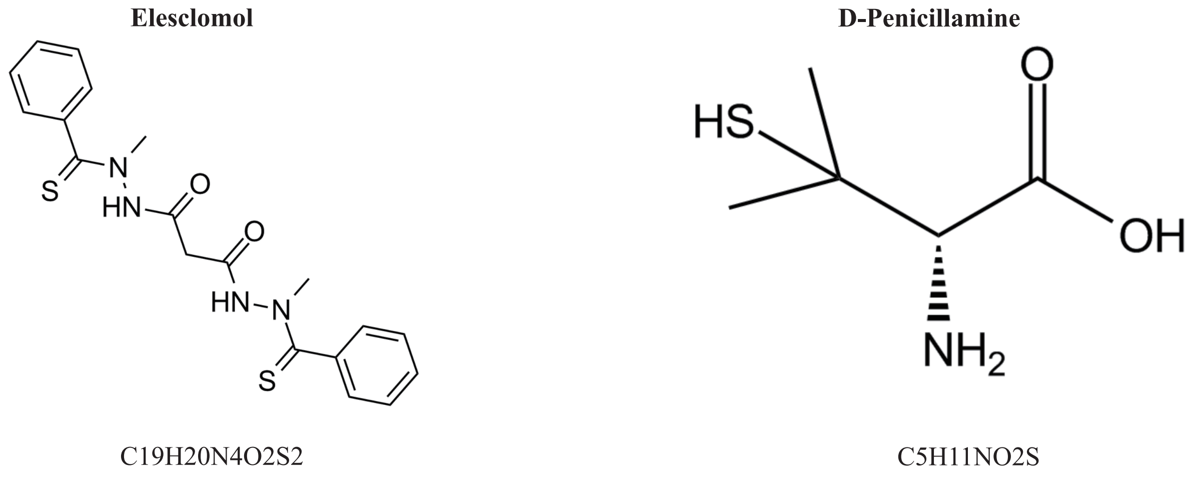


**Figure S4 Compound structures for Elesclomol and D-Penicillamine in the study**

### Source Western-blot Images for Figure 11C

#### Repeat 1

β-tubulin

SLC31A1

FDX1

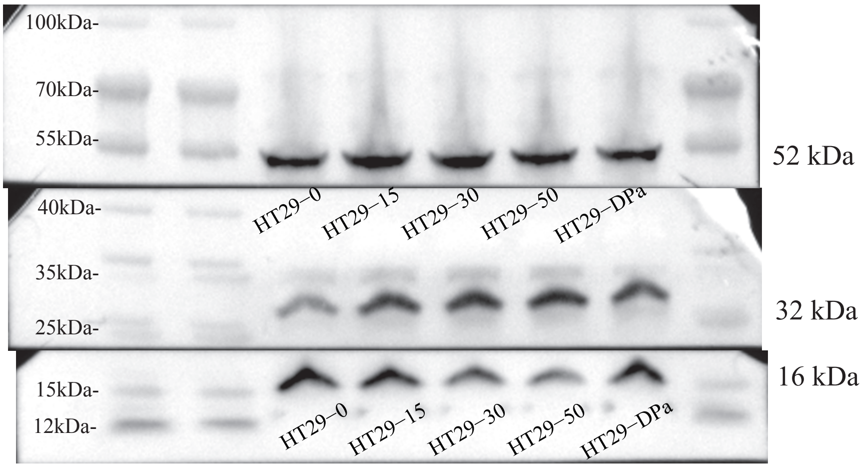


#### Repeat 2

β-tubulin

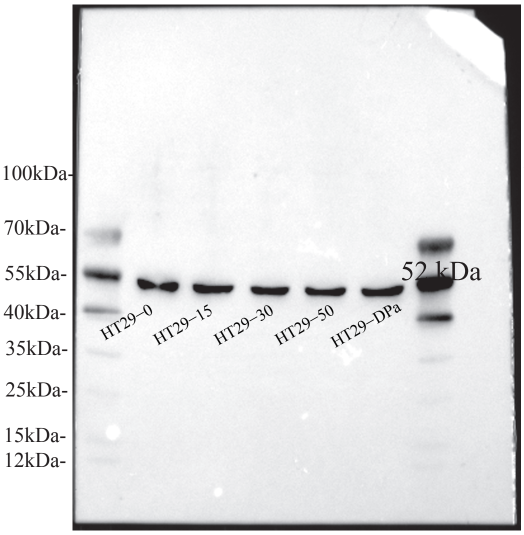


SLC31A1

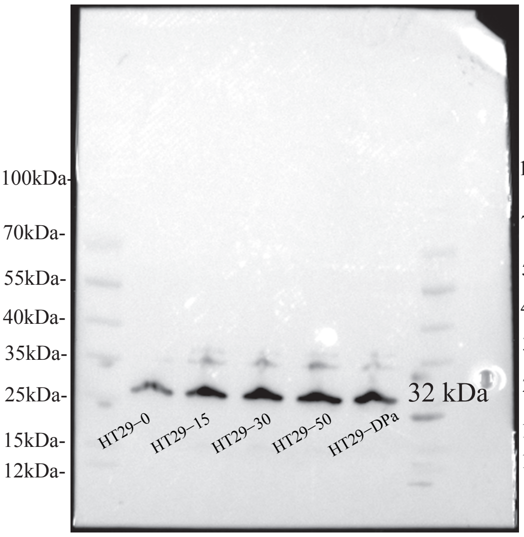


FDX1

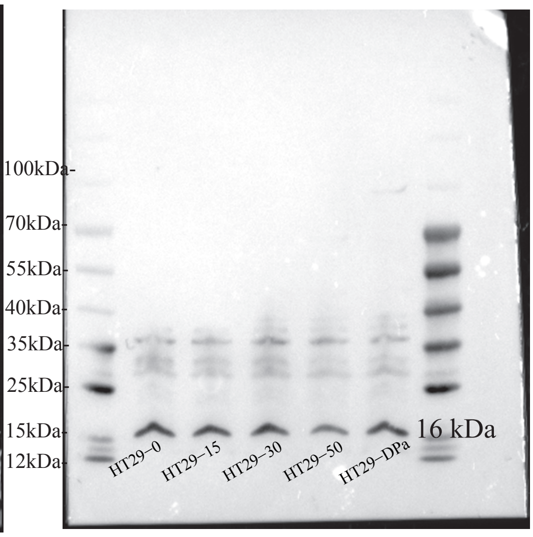


#### Repeat 3

β-tubulin

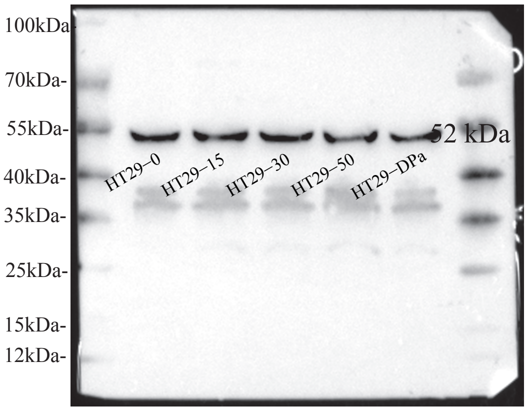


SLC31A1

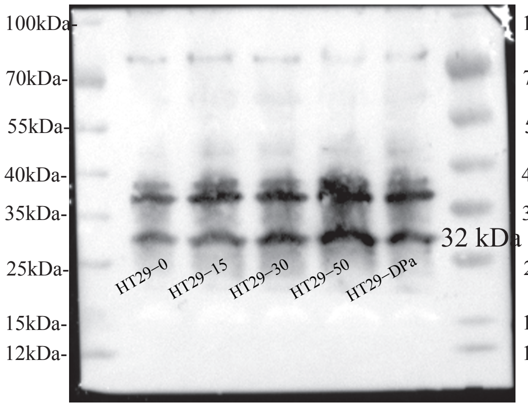


FDX1

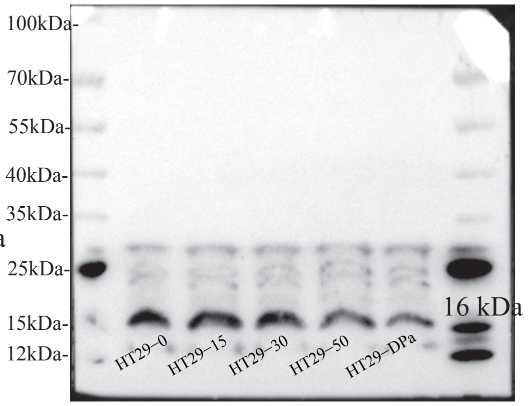


**FigureS5 Source Western-blot Images for Figure 11C**

### Source Western-blot Images for Figure 11D

#### Repeat 1

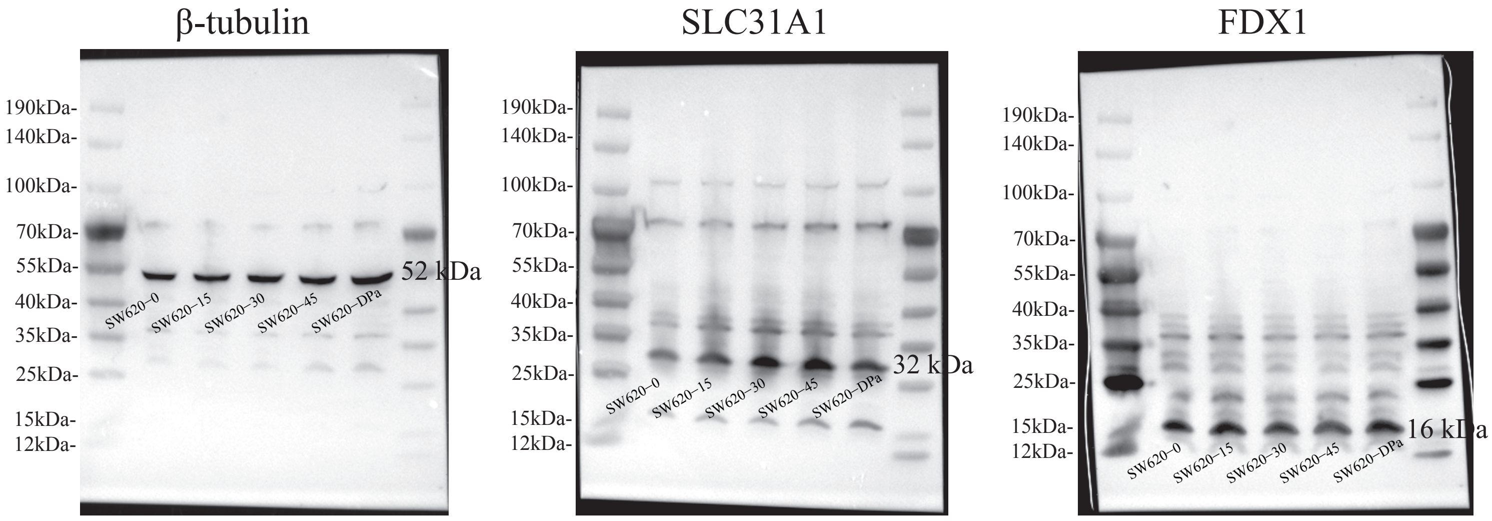


#### Repeat 2

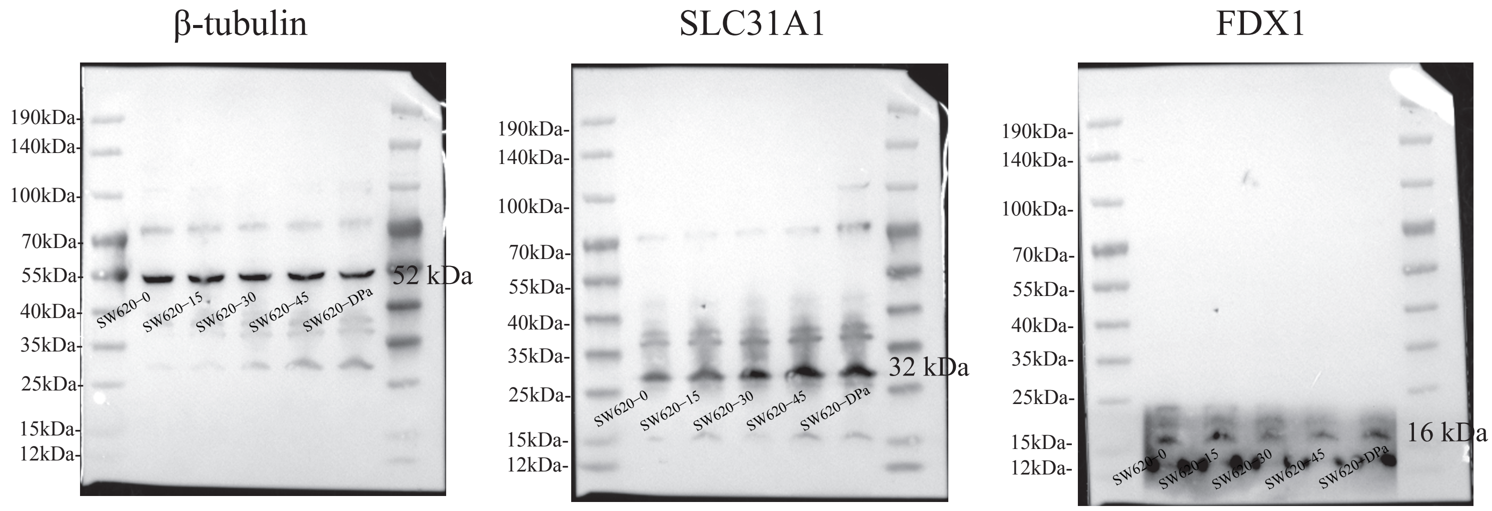


#### Repeat 3

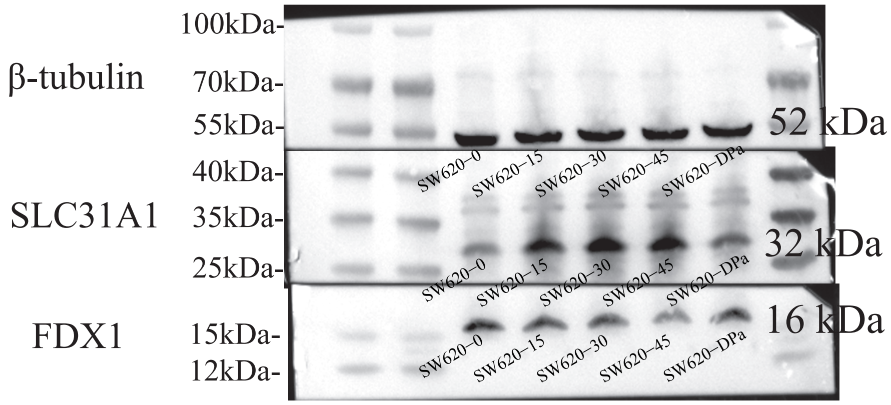


**FigureS6 Source Western-blot Images for Figure 11D**

**Supplementary Tables**

**Table S1 The full name, function of the 16 CPRMs**

| Gene | Full Name | Function |
|---|---|---|
| FDX1 | Ferredoxin 1 | Transfers electrons from NADPH through ferredoxin reductase to mitochondrial cytochrome P450, involved in steroid, vitamin D, and bile acid metabolism. |
| LIPT1 | Lipoyltransferase 1 | Transfers the lipoyl moiety to apoproteins which is the second step of the process of transferring lipoic acid to proteins. |
| LIAS | Lipoic Acid Synthetase | Catalyzes the final step in the de novo pathway for the biosynthesis of lipoic acid, a potent antioxidant. |
| DLD | Dihydrolipoamide Dehydrogenase | The encoded protein has been identified as a moonlighting protein based on its ability to perform mechanistically distinct functions. |
| DBT | Dihydrolipoamide Branched Chain Transacylase E2 | This gene encodes the transacylase (E2) subunit of branched-chain alpha-keto acid dehydrogenase complex (BCKD) which is an inner-mitochondrial enzyme complex involved in the breakdown of the branched-chain amino acids isoleucine, leucine, and valine. |
| GCSH | Glycine Cleavage System Protein H | The protein encoded by this gene is the H protein, a component of glycine cleavage system, which transfers the methylamine group of glycine from the P protein to the T protein |
| DLST | Dihydrolipoamide S-succinyltransferase | This gene encodes a mitochondrial protein which is one of the three components (the E2 component) of the 2-oxoglutarate dehydrogenase complex that catalyzes the overall conversion of 2-oxoglutarate to succinyl-CoA and CO(2). |
| DLAT | Dihydrolipoamide S-acetyltransferase | This gene encodes component E2 of the multi-enzyme pyruvate dehydrogenase complex (PDC). The protein product of this gene, dihydrolipoamide acetyltransferase, accepts acetyl groups formed by the oxidative decarboxylation of pyruvate and transfers them to coenzyme A. |
| PDHA1 | Pyruvate Dehydrogenase E1 Subunit Alpha 1 | This gene encodes the E1 alpha 1 subunit containing the E1 active site, and plays a key role in the function of the PDH complex which provides the primary link between glycolysis and the tricarboxylic acid (TCA) cycle. |
| PDHB | Pyruvate Dehydrogenase E1 Subunit Beta | This gene encodes beta subunit of the E1 enzyme, which is a component of the pyruvate dehydrogenase (PDH) complex. |

| | | |
|---|---|---|
| SLC31A1 | Solute Carrier Family 31 Member 1 | The protein encoded by this gene is a high-affinity copper transporter found in the cell membrane. |
| ATP7A | ATPase Copper Transporting Alpha | This gene encodes a transmembrane protein that functions in copper transport across membranes. |
| ATP7B | ATPase Copper Transporting Beta | This protein is a monomer, and functions as a copper-transporting ATPase which exports copper out of the cells, such as the efflux of hepatic copper into the bile. |
| CDKN2A | Cyclin Dependent Kinase Inhibitor 2A | At least three alternatively spliced variants encoding distinct proteins have been reported, two of which encode structurally related isoforms known to function as inhibitors of CDK4 kinase. |
| GLS | Glutaminase | This gene encodes the K-type mitochondrial glutaminase. The encoded protein is an phosphate-activated amidohydrolase that catalyzes the hydrolysis of glutamine to glutamate and ammonia. |
| MTF1 | Metal Regulatory Transcription Factor 1 | This gene encodes a transcription factor that induces expression of metallothioneins and other genes involved in metal homeostasis in response to heavy metals such as cadmium, zinc, copper, and silver. |

**Table S2 The clinical characteristics of the individuals in our transcriptome cohort**

| Patients | Age | Gender | Vessel embolus | Nerve invasion | Cancerous node | EGFR | pTNM | TNM Stage | CuproScore | Group |
|---|---|---|---|---|---|---|---|---|---|---|
| 1 | 69 | male | (-) | (-) | (-) | (±) | T3N0M0 | II | 0.692436901 | Low |
| 2 | 58 | male | (-) | (-) | (-) | (+) | T3N2aM1 | IV | 2.0065503 | High |
| 3 | 68 | male | (-) | (-) | (-) | (-) | T3N0M0 | II | 0.97281857 | High |
| 4 | 57 | male | (-) | (-) | (-) | (+) | T3N0M0 | II | 1.493446474 | High |
| 5 | 56 | female | (-) | (+) | (+) | (+) | T3N0M0 | II | -1.45929755 | Low |
| 6 | 38 | male | (-) | (-) | (-) | (+) | T3N0M0 | II | 0.921750987 | Low |
| 7 | 56 | male | (-) | (-) | (-) | (+) | T3N0M0 | II | -11.7853259 | Low |
| 8 | 50 | male | (-) | (-) | (-) | (+) | T3N0M0 | II | 0.490919602 | Low |
| 9 | 66 | male | (-) | (-) | (-) | (+) | T3N0M0 | II | 2.078366947 | High |
| 10 | 56 | male | (-) | (-) | (-) | (+) | T3N1M0 | III | 1.654392402 | High |
| 11 | 47 | female | (+) | (-) | (-) | (+) | T2N2bM0 | III | 0.109704117 | Low |
| 12 | 72 | female | (-) | (-) | (-) | (±) | T3N1M0 | III | 2.824237097 | High |

**Table S3 Real-time PCR primer sequences of 16 CPRMs**

| Name | Description | Primer Sequences（5’ to 3’） |
|---|---|---|
| ATP7B(F) | Forward Primer | GGCCGTCATCACTTATCAGCC |
| ATP7B(R) | Reverse Primer | GGGAGCCACTTTGCTCTTGA |
| SLC31A1(F) | Forward Primer | GGGGATGAGCTATATGGACTCC |
| SLC31A1(R) | Reverse Primer | TCACCAAACCGGAAAACAGTAG |
| DLST(F) | Forward Primer | GAACTGCCCTCTAGGGAGAC |
| DLST(R) | Reverse Primer | AACCTTCCTGCTGTTAGGGTA |
| GCSH(F) | Forward Primer | GGAAGCGTTGGGAGATGTTGT |
| GCSH(R) | Reverse Primer | TCTGAAGGGTTACTCAGTGTCA |
| DBT(F) | Forward Primer | CAGTTCGCCGTCTGGCAAT |
| DBT(R) | Reverse Primer | CCTGTGAATACCGGAGGTTTTG |
| ATP7A(F) | Forward Primer | TGACCCTAAACTACAGACTCCAA |
| ATP7A(R) | Reverse Primer | CGCCGTAACAGTCAGAAACAA |
| FDX1(F) | Forward Primer | GCCTCTTTGGAGTCTCTCGC |
| FDX1(R) | Reverse Primer | CCCAACCGTGATCTGTCTGT |
| LIAS(F) | Forward Primer | CAGCCCAGTCAGACCGTTAAG |
| LIAS(R) | Reverse Primer | TTTCTGGCGTTTTAGGTTTCCT |
| LIPT1(F) | Forward Primer | CCTCTGTTGTAATTGGTAGGCAT |
| LIPT1(R) | Reverse Primer | CTGGGGTTGGACAGCATTCAG |
| DLD(F) | Forward Primer | CTCATGGCCTACAGGGACTTT |
| DLD(R) | Reverse Primer | GCATGTTCCACCAAGTGTTTCAT |
| DLAT(F) | Forward Primer | CGGAACTCCACGAGTGACC |
| DLAT(R) | Reverse Primer | CCCCGCCATACCCTGTAGT |
| PDHA1(F) | Forward Primer | TGGTAGCATCCCGTAATTTTGC |
| PDHA1(R) | Reverse Primer | ATTCGGCGTACAGTCTGCATC |
| PDHB(F) | Forward Primer | AAGAGGCGCTTTCACTGGAC |
| PDHB(R) | Reverse Primer | ACTAACCTTGTATGCCCCATCA |
| MTF1(F) | Forward Primer | CACAGTCCAGACAACAACATCA |
| MTF1(R) | Reverse Primer | GCACCAGTCCGTTTTTATCCAC |
| GLS(F) | Forward Primer | TCTACAGGATTGCGAACGTCT |
| GLS(R) | Reverse Primer | CTTTGTCTAGCATGACACCATCT |
| CDKN2A(F) | Forward Primer | GATCCAGGTGGGTAGAAGGTC |
| CDKN2A(R) | Reverse Primer | CCCCTGCAAACTTCGTCCT |
